# Computational exploration of the global microbiome for antibiotic discovery

**DOI:** 10.1101/2023.08.31.555663

**Authors:** Célio Dias Santos-Júnior, Marcelo Der Torossian Torres, Yiqian Duan, Álvaro Rodríguez del Río, Thomas S.B. Schmidt, Hui Chong, Anthony Fullam, Kuhn Michael, Chengkai Zhu, Amy Houseman, Jelena Somborski, Anna Vines, Xing-Ming Zhao, Peer Bork, Jaime Huerta-Cepas, Cesar de la Fuente-Nunez, Luis Pedro Coelho

**Affiliations:** Institute of Science and Technology for Brain-Inspired Intelligence - ISTBI, Fudan University, Shanghai, China; Machine Biology Group, Departments of Psychiatry and Microbiology, Institute for Biomedical Informatics, Institute for Translational Medicine and Therapeutics, Perelman School of Medicine, University of Pennsylvania; Philadelphia, Pennsylvania, United States of America; Departments of Bioengineering and Chemical and Biomolecular Engineering, School of Engineering and Applied Science, University of Pennsylvania; Philadelphia, Pennsylvania, United States of America; Penn Institute for Computational Science, University of Pennsylvania; Philadelphia, Pennsylvania, United States of America; Centro de Biotecnología y Genómica de Plantas, Universidad Politécnica de Madrid (UPM) - Instituto Nacional de Investigación y Tecnología Agraria y Alimentaria (INIA-CSIC), Campus de Montegancedo-UPM, 28223 Pozuelo de Alarcón, Madrid, Spain; Structural and Computational Biology Unit, European Molecular Biology Laboratory, Heidelberg, Germany; Max Delbrück Centre for Molecular Medicine, Berlin, Germany; Department of Bioinformatics, Biocenter, University of Würzburg, Würzburg, Germany; Department of Neurology, Zhongshan Hospital, Fudan University, Shanghai, China; State Key Laboratory of Medical Neurobiology, Institutes of Brain Science, Fudan University, Shanghai, China; MOE Key Laboratory of Computational Neuroscience and Brain-Inspired Intelligence; MOE Frontiers Center for Brain Science, Fudan University, Shanghai, China; International Human Phenome Institute, Shanghai, China

**Keywords:** metagenomics, antimicrobial peptides, antimicrobial activity, machine learning, global microbiome

## Abstract

Novel antibiotics are urgently needed to combat the antibiotic-resistance crisis. We present a machine learning-based approach to predict prokaryotic antimicrobial peptides (AMPs) by leveraging a vast dataset of 63,410 metagenomes and 87,920 microbial genomes. This led to the creation of AMPSphere, a comprehensive catalog comprising 863,498 non-redundant peptides, the majority of which were previously unknown. We observed that AMP production varies by habitat, with animal-associated samples displaying the highest proportion of AMPs compared to other habitats. Furthermore, within different human-associated microbiota, strain-level differences were evident. To validate our predictions, we synthesized and experimentally tested 50 AMPs, demonstrating their efficacy against clinically relevant drug-resistant pathogens both in vitro and in vivo. These AMPs exhibited antibacterial activity by targeting the bacterial membrane. Additionally, AMPSphere provides valuable insights into the evolutionary origins of peptides. In conclusion, our approach identified AMP sequences within prokaryotic microbiomes, opening up new avenues for the discovery of antibiotics.

## Introduction

Antibiotic-resistant infections are becoming increasingly difficult to treat with conventional therapies. Indeed, such infections currently kill 1.27 million people per year^1^. Therefore, there is an urgent need to develop novel methods for antibiotic discovery. Computational approaches have been recently developed to accelerate our ability to identify novel antibiotics, including antimicrobial peptides (AMPs)^2–5^.

AMPs, found in all domains of life^6–9^, are short sequences (here operationally defined as 10- 100 amino acid residues^10^) capable of disturbing microbial growth^7,11^. AMPs can target proteins, RNA, DNA, and other intracellular molecules; they most commonly interfere with cell wall integrity and cause cell lysis^7^. Natural AMPs can originate by proteolysis^3,12^ or by non-ribosomal synthesis^13^, or they can be encoded within the genome^14^. Recently, proteome mining approaches have been developed to identify antimicrobials in extinct organisms^15^.

Bacteria live in an intricate balance of antagonism and mutualism in natural habitats. AMPs play an important role in modulating such microbial interactions and can displace competitor strains, facilitating cooperation^16^. For instance, pathogens such as *Shigella* spp.^17^, *Staphylococcus* spp.^18^, *Vibrio cholerae*^19^, and *Listeria* spp.^20,21^ produce AMPs that eliminate competitors (sometimes from the same species) and occupy their niche.

AMPs hold promise as potential therapeutics and have already been used clinically as antiviral drugs (*e.g.*, enfuvirtide and telaprevir^22^). AMPs that exhibit immunomodulatory properties are currently undergoing clinical trials^23^, as are AMPs that may be used to address yeast and bacterial infections^24^ (*e.g.*, pexiganan, LL-37, PAC-113). Although most AMPs display broad-spectrum activity, some can present narrow activity, having activity only against closely related members of the same species or genus^25^. Such AMPs are more targeted agents than conventional broad-spectrum antibiotics^26,27^. For example, the FDA-approved small peptide nisin has been shown to restore microbiome homeostasis in *Clostridioides difficile* infections in mice^28^ and in *ex vivo* models of the human gut^29^. Furthermore, contrary to conventional antibiotics, the evolution of resistance to many AMPs occurs at low rates and is not related to cross-resistance to other classes of widely used antibiotics^3,30,31^.

The application of metagenomic analyses to the study of AMPs has been limited due to technical constraints, primarily stemming from the challenge of distinguishing genuine protein-coding sequences from false positives^32^. As a consequence, the significance of small open reading frames (smORFs) has been historically overlooked in (meta)genomic analyses^33–35^. In recent years, significant progress has been made in metagenomic analyses of human-associated small open reading frames (smORFs)^5,36^. These advancements have incorporated machine learning (ML) techniques to identify smORFs encoding proteins belonging to specific functional categories^37–40^. Notably, a recent study uncovered approximately 2,000 AMPs from metagenomic samples of human gut microbiomes^5^. Nevertheless, it is important to note that the human gut represents only a fraction of the overall microbial diversity, suggesting that there remains an immense potential for the discovery of AMPs from prokaryotes in the diverse range of habitats across the globe.

In this study, we employed ML to predict and catalog the entire global microbiome. By computationally exploring 63,410 publicly available metagenomes and 87,920 high-quality microbial genomes^41^, we uncovered a vast array of AMP diversity. This resulted in the creation of AMPSphere, a collection of 863,498 non-redundant peptide sequences, encompassing candidate AMPs (c_AMPs) derived from (meta)genomic data. Remarkably, the majority of these identified c_AMP sequences had not been previously described. Our analysis revealed that these AMPs were specific to particular habitats and were predominantly not core genes in the pangenome.

Moreover, we synthesized 50 c_AMPs from AMPSphere and found that 54% of them exhibited antimicrobial activity *in vitro* against clinically significant ESKAPEE pathogens, which are recognized as public health concerns^42,43^. These peptides demonstrated their ability to target bacterial membranes and were prone to adopting α-helical and β-structures. Notably, the leading candidates displayed promising anti-infective activity in a preclinical animal model.

## Results

### AMPSphere comprises almost 1 million c_AMPs from several habitats

AMPSphere incorporates c_AMPs predicted with ML using Macrel^40^ from 63,410 worldwide distributed publicly available metagenomes (***Fig. 1A*** and ***Table SI1***) and 87,920 high-quality bacterial and archaeal genomes^41^. Sequences present in a single sample were removed^40^, except when they had a significant match (defined as amino acids identity ≥ 75% and E-value ≤ 10⁻⁵) to a sequence in DRAMP 3.0^44^. This resulted in 5,518,294 genes, 0.1% of the total predicted smORFs, coding for 863,498 non-redundant c_AMPs (on average 37±8 residues long; ***Fig. 1A*** and ***SI1***).

**Figure 1.**
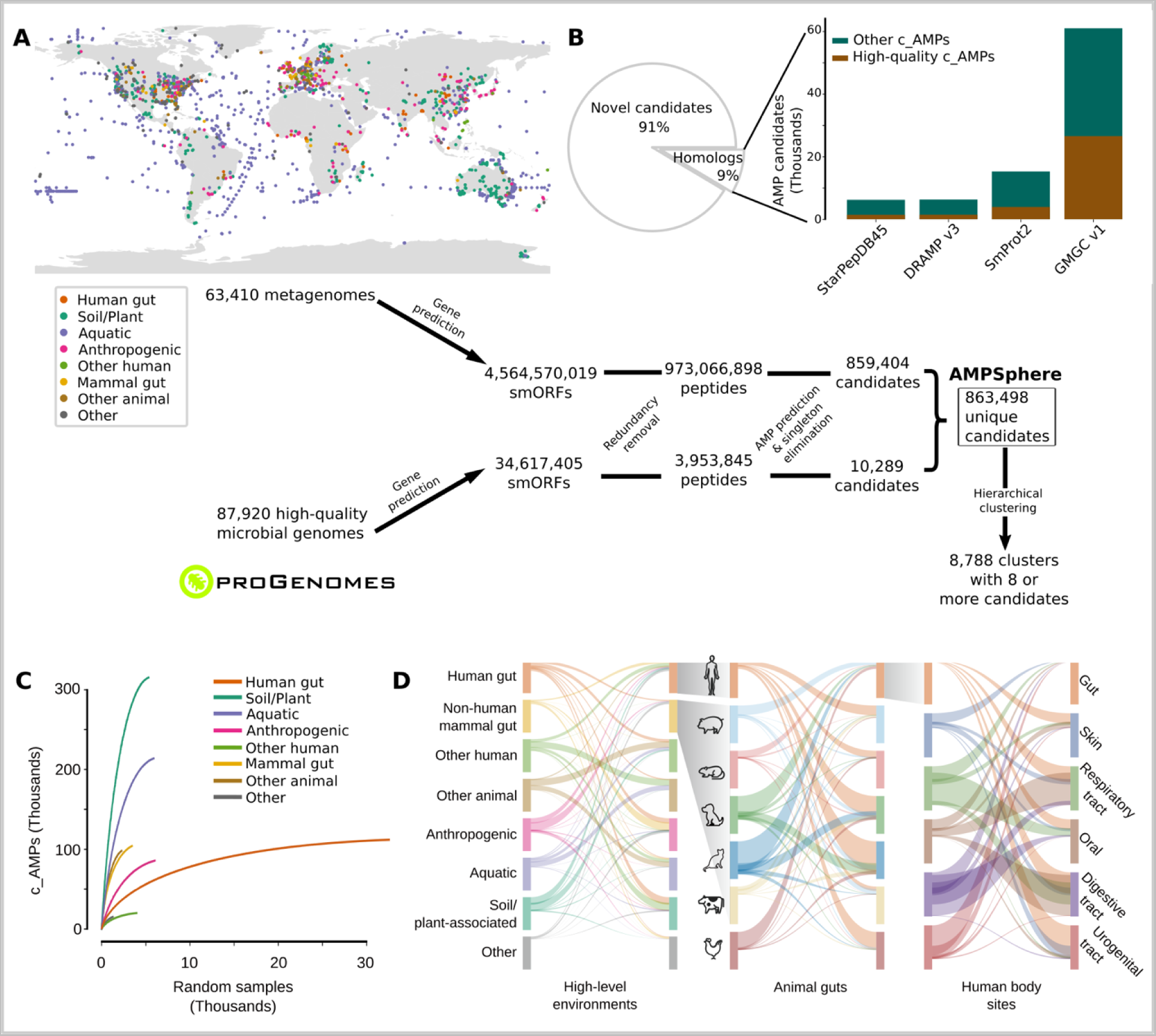
AMPSphere comprises 836,498 non-redundant c_AMPs from thousands of metagenomes and high-quality microbial genomes. (***A***) To build AMPSphere, we first assembled 63,410 publicly available metagenomes from diverse habitats. A modified version of Prodigal^32^, which can also predict smORFs (30-300 bp), was used to predict genes on the resulting metagenomic contigs as well as on 87,920 microbial genomes from ProGenomes2^41^. Macrel^40^ was applied to the 4,599,187,424 predicted smORFs to obtain 863,498 non-redundant c_AMPs (see also ***Fig. SI1***). c_AMPs were then hierarchically clustered in a reduced amino acids alphabet using 100%, 85%, and 75% identity cutoffs. We observed at 75% of identity 118,051 non-singleton clusters, and 8,788 of them were considered families (≥ 8 c_AMPs). (***B***) Only 9% of c_AMPs have detectable homologs in other peptides (SmProt 2^45^, DRAMP 3.0^44^, starPepDB 45k^91^, STsORFs^113^) and general protein datasets (GMGCv1^46^) - see also ***Fig. SI2B.*** (***C***) AMP discovery is impacted by the sampling, with most of the habitats presenting steep sampling curves, *e.g.*, soil. (***D***) Overall, c_AMPs are habitat-specific - see also ***Fig. SI2C-D*** and ***Tables SI1 and SI2***.

We subsequently estimated the quality of the smORF predictions and detected 20% (172,840) of the c_AMP sequences in independent metaproteomes or metatranscriptomes (***Fig. SI2A*** and ***Methods - Quality control of c_AMPs***). To further assess the gene predictions, we subjected the peptides to a bundle of *in silico* quality tests (see ***Methods - Quality control of c_AMPs***) and the subset of c_AMPs that passed all of them (9.2%) is hereafter designated as high-quality.

Only 0.7% of the identified c_AMPs (6,339 peptides) are homologous to experimentally validated AMP sequences in the DRAMP 3.0 database^44^. Moreover, most c_AMPs were also absent from protein databases not specific to AMPs (***Fig. 1B***), such as SmProt 2^45^ or GMGCv1^46^, suggesting that the c_AMPs represent an entirely novel region of peptide sequence space. In total, we could find only 73,774 (8.5%) c_AMPs with homologs in the tested databases. High-quality c_AMPs were detected in public databases with higher frequency than general c_AMPs (2.5-fold, P_Hypergom._ = 4.2·10^−250^, ***Fig. 1B***).

To put c_AMPs in an evolutionary context, we hierarchically clustered peptides using a reduced amino acid alphabet of 8 letters^47^ and identity cutoffs of 100%, 85%, and 75% (***Fig. SI3***). At the 75% identity level, we obtained 521,760 protein clusters of which 405,547 were singletons, corresponding to 47% of all c_AMPs from AMPSphere. A total of 78,481 (19.3%) of these singletons were detected in metatranscriptomes or metaproteomes from various sources, indicating that they are not artifacts. The large number of singletons suggests that most c_AMPs originated from processes other than diversification within families, which is the opposite of the supposed origin of full-length proteins, in which singleton families are rare^46^. The 8,788 clusters with ≥8 peptides obtained at 75% of identity are hereafter named “families’’, as in Sberro et al.^36^. Among them, we consider 6,499 as high-quality families because they contain evidence of translation or transcription, or ≥75% of their sequences pass all *in silico* quality tests (see Methods). These high-quality families span 15.4% of the AMPSphere (133,309 peptides).

All the c_AMPs predicted here can be accessed at https://ampsphere.big-data-biology.org/. Users can retrieve the peptide sequences, ORFs, and predicted biochemical properties of each c_AMP (*e.g.*, molecular weight, isoelectric point, and charge at pH 7.0). We also provide the distribution across geographical regions, habitats, and microbial species for each c_AMP.

### c_AMPs are rare and habitat specific

The AMPSphere spans 72 different habitats, which were classified into 8 high-level habitat groups, *e.g*., soil/plant (36.6% of c_AMPs in AMPSphere), aquatic (24.8%), human gut (13%) - (***Fig. 1A*** and ***Table SI2)***. Most of the habitats, except for the human gut, appear to be far from saturation in terms of newly discovered c_AMPs (***Fig. 1C***). In fact, most AMPs are rare (median number of detections is 99, or 0.17% of the dataset), with 83.97% being observed in <1% of samples - see ***Fig. SI2***. Only 10.8% (93,280) c_AMPs were detected in more than one high-level habitat (henceforth, “multi-habitat c_AMPs”); this level is 7.25-fold less frequent than would be expected by a random assignment of habitats to samples (P_Permutation_ < 10^−300^, see ***Methods – Multi-habitat and rare c_AMPs***). Even within high-level habitat groups, c_AMP contents overlap between habitats much less than expected by chance (2.4 to 192-fold less, P_Permutation_⩽ 5.4·10^−50^, see ***Methods - Significance of the overlap of c_AMP contents***; ***Fig. 1D***), indicating the existence of cross-habitat boundaries.

### Mutations in larger genes generate c_AMPs as independent genomic entities

Many AMPs are generated post-translationally by the fragmentation of larger proteins^12^. In parallel, encrypted peptides from protein sequences within the human proteome were also shown to be highly active. Those peptides have different physicochemical features compared to known AMPs^3,31^. However, AMPSphere only considered peptides encoded by dedicated genes. Nonetheless, we hypothesized that some of these have originated from larger proteins by fragmentation at the genomic level: about 7% (61,020) of AMPSphere c_AMPs are homologous to full-length proteins in GMGCv1^46^ (***Fig. 1B)***, with 27% of hits sharing the start codon with the full-length protein, which suggests early termination of full-length proteins as one mechanism for generating novel c_AMPs (***Fig. 2A*** and ***2B***).

**Figure 2.**
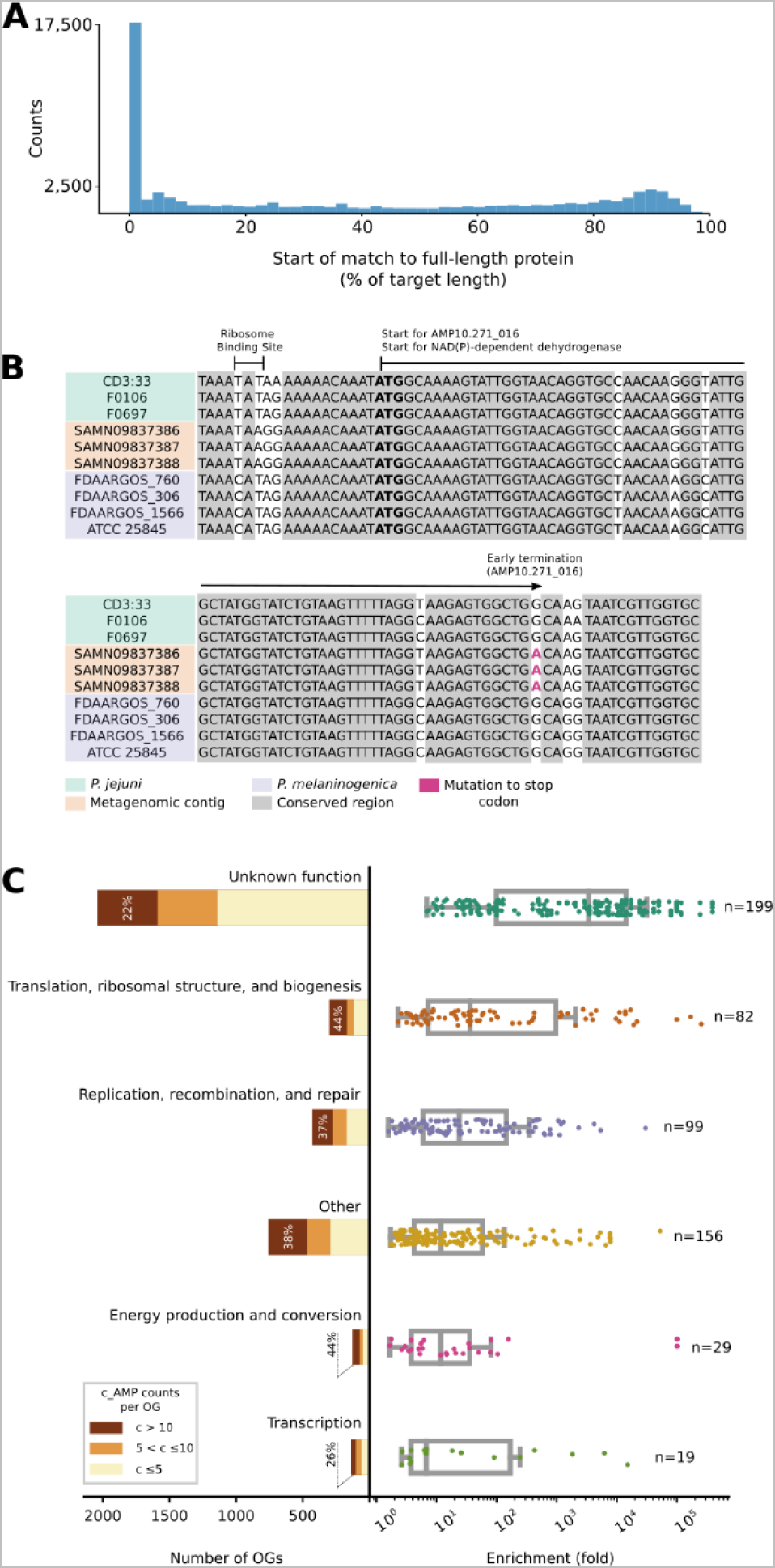
Mutations in genes encoding large proteins generate c_AMPs as independent genomic entities. (***A***) About 7% of c_AMPs are homologous to proteins from GMGCv1^46^, with almost one-fourth of the hits sharing start positions with the larger protein. (***B***) As an illustrative example, AMP10.271_016 was recovered in three samples of human saliva from the same donor^132^. AMP10.271_016 is predicted to be produced by *Prevotella jejuni*, sharing the start codon (bolded) of an NAD(P)-dependent dehydrogenase gene (WP_089365220.1), the transcription of which was stopped by a mutation (in red; TGG > TGA). (***C***) The OGs of unknown function represent the largest (2,041 out of 3,792 OGs) and most enriched (P_Kruskal_ = 2.66·10^−39^) class with homologs to c_AMPs in GMGCv1^46^. Interestingly, when considered individually, the number of c_AMP hits to unknown OGs was the lowest (P_Kruskal_ = 6·10^−3^). These results do not change when underrepresented OGs are excluded by using different thresholds (*e.g*., at least 10, 20, or 100 homologs per OG) - see also ***Table SI3***.

To investigate the function of the full-length proteins homologous to AMPs, we mapped the matching proteins from GMGCv1^46^ to their orthologous groups (OGs) from eggNOG 5.0^48^. We identified 3,792 (out of 43,789) OGs significantly enriched (P_Hypergeom._ < 0.05, after multiple hypothesis corrections with the Holm-Sidak method) among the hits from AMPSphere. Although OGs of unknown function comprise 53.8% of all identified OGs, when considered individually, these OGs are, on average, smaller than OGs in other categories. Thus, despite each OG having a relatively small number of c_AMP hits, when compared to the background distribution of the OGs in GMGCv1, OGs of unknown function were the most enriched among the c_AMP hits, with an average enrichment of 10,857 fold (P_Mann_ ≤ 3.9·10^−4^; ***Fig. 2C***; ***Table SI3***).

### c_AMP genes may arise after gene duplication events

We next raised the question of whether c_AMPs would be predominantly present in specific genomic contexts. To investigate the functions of the neighboring genes of the c_AMPs, we mapped them against 169,484 high-quality genomes included in a study by del Río et al.^49^. 38.9% (21,465 out of 55,191) c_AMPs with more than two homologs in different genomes in the database show phylogenetically conserved genomic context with genes of known function (see ***Methods - Genomic context conservation analysis***). This proportion of c_AMPs in conserved genomic context is slightly higher than for other gene family clusters calculated on the genomes’ gene sets (1.1-fold, P_Permutation_ = 1.3·10^−111^). This difference becomes more pronounced when comparing the genomic context of c_AMPs and protein families composed of short (< 50 amino acids) peptides (11-fold increase, P_Permutation_ < 10^−330^).

Despite being involved in similar processes, AMPs were significantly depleted from conserved genomic contexts involving known systems of antibiotic synthesis and resistance, even when they were compared to small protein families (0.6-fold, P_Permutation_ = 1.7·10^−8^, ***Fig. 3***). Instead, we found that c_AMPs are encoded in conserved genomic contexts with ribosomal genes (24.1%) and ABC transporters (18%) at a higher frequency than other gene families (***Fig. 3A*** and ***Tables SI4*** and ***SI5***).

**Figure 3.**
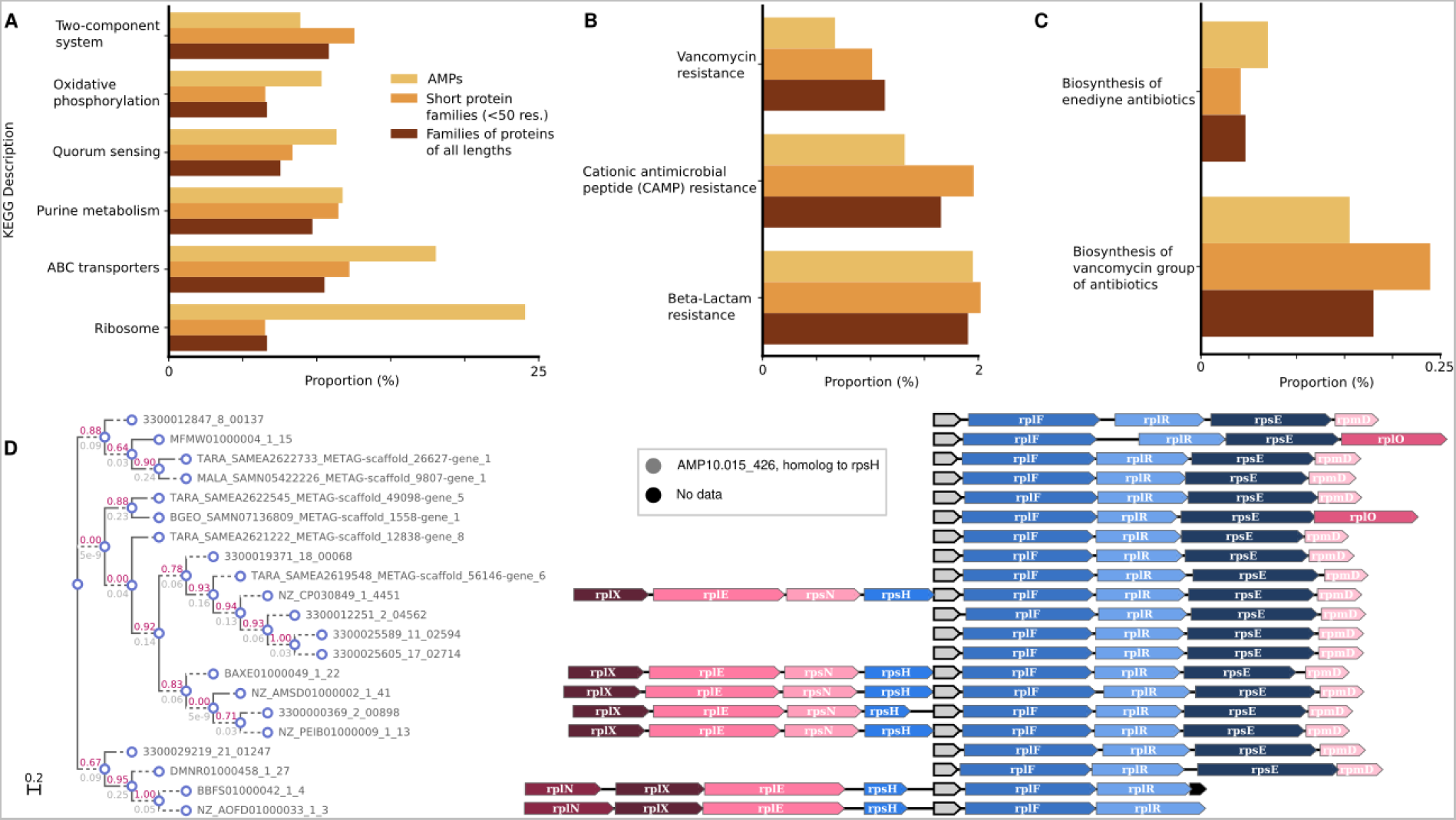
*The genome context of c_AMPs shows a preference for neighborhoods containing ABC transporters and ribosome assembly proteins* - see *Tables SI4* and *SI5.* **(*A*)** Compared to other proteins, c_AMPs tend to be closer to ABC transporters and ribosomal machinery-related genes than families of proteins with different sizes (≤ 50 amino acids and all lengths). **(*B*)** The proportion of c_AMPs in a genome context involving antibiotic resistance genes is lower than families of proteins shorter than 50 amino acids and, in the case of CAMP and vancomycin resistance, than all proteins. **(*C*)** The proportion of c_AMPs in neighborhoods with antibiotic synthesis-related genes is very small (<0.25%). **(*D*)** AMP10.015_426 is an example of a c_AMP homologous to the ribosomal protein rpsH, found in the context of other ribosomal protein genes.

Most of the c_AMPs (2,201 out of the 2,642) in conserved ribosomal genomic contexts are homologous to ribosomal proteins (***Fig. 3D***), congruent with the observation that, in some species, ribosomal proteins have antimicrobial properties^50^. Seventy-seven of the c_AMPs that were homologous to ribosomal proteins were homologous to a gene in their immediate vicinity (up to 1 gene up/downstream). This phenomenon is not exclusive to ribosomal proteins: 2,309 c_AMPs can be annotated to the same KEGG Orthologous Group (KO) as some of their conserved neighbors and may have originated from gene duplication events, the common annotation being interpreted, in this context, as evidence for a common evolutionary origin and not as a functional prediction for the c_AMPs. Interestingly, 1,707 (73.9%) of these c_AMPs are located downstream of the conserved neighbor with the same KO annotation. The livM family (branched-chain amino acid transport system permease protein, K01998), a transposase family (K07486), and a class of permeases (K03106) are the most common KOs assigned to c_AMPs and their neighbors (185, 128, and 67 c_AMPs, respectively) - see ***Table SI6***.

### Most c_AMPs are members of the accessory pangenome

We observed that only a small portion (5.9%, P_Permutation_ = 4.8·10^−3^, N_Species_ = 416) of c_AMP families present in ProGenomes2^41^ show prevalence ≥95% in genomes from the same species (***Fig. 4***), here referred to as “core”^51^. This is consistent with previous work, in which AMP production was observed to be strain-specific^52^. In contrast, a high proportion (*circa* 68.8%) of full-length protein families are core in ProGenomes2^41^ species. There is a 1.9-fold greater chance (P_Fisher_ = 2.2·10^−92^) of finding a pair of genomes from the same species sharing at least one c_AMP when they belong to the same strain (99.5% ≤ ANI < 99.99%).

**Figure 4.**
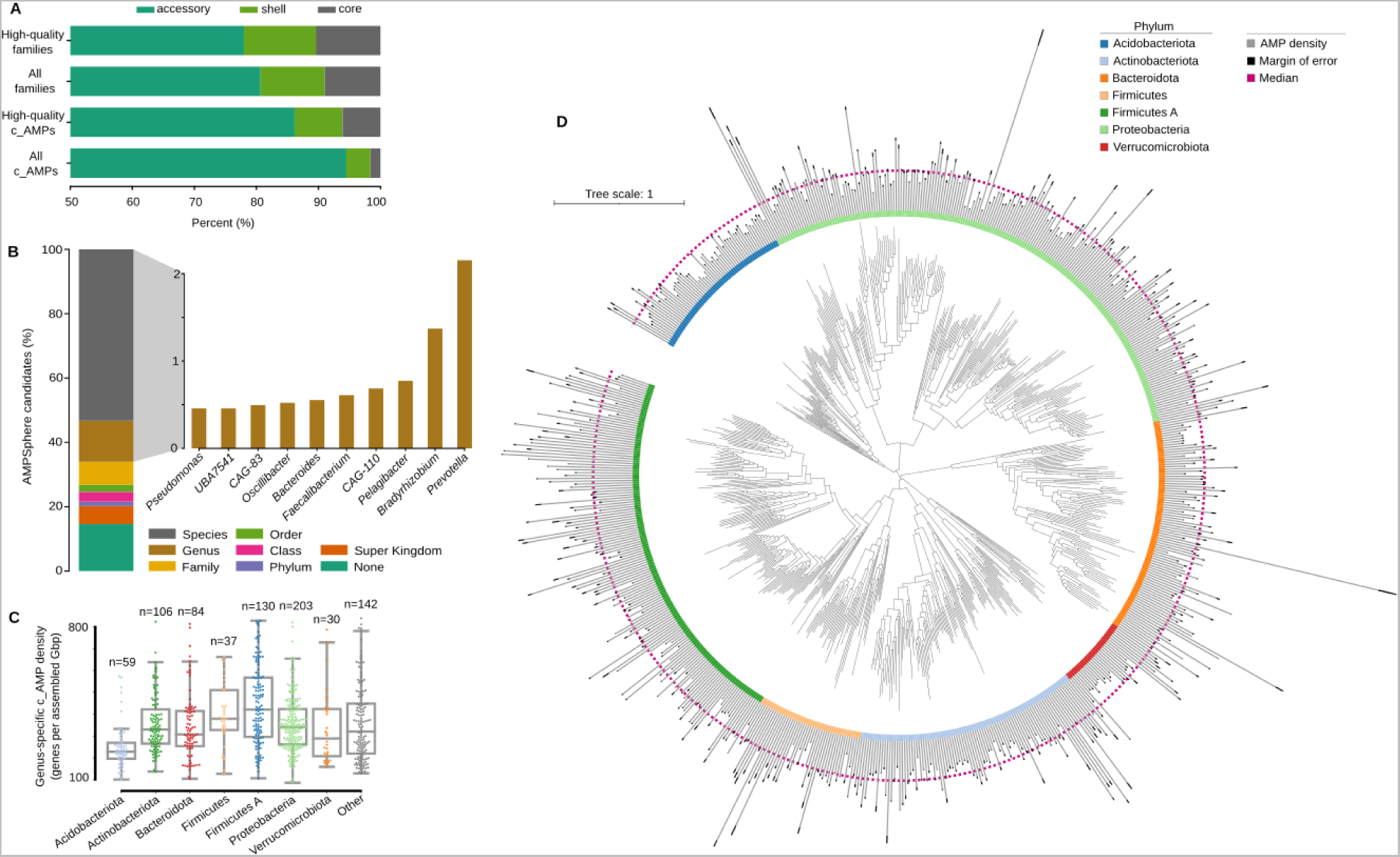
AMP variation is taxonomy-dependent. (***A***) Most of c_AMPs and families from ProGenomes2^41^ are classified as core genes (defined as ≥95% genomes within a species). (***B***) The majority of the c_AMPs were classified down to the level of genus and species. Animal-associated genera (*e.g. Prevotella*, *Faecalibacterium*, *CAG-110*) contribute the most c_AMPs, possibly reflecting data sampling. (***C***) Using the ⍴*_AMP_* per genus, we observed the distribution of c_AMPs per phyla, with Bacillota A as the densest. (***D***) The ⍴*_AMP_* distribution (gray bars, confidence interval of 95% shown as black bars) with respect to taxonomy shows Bacillota A, Actinomycetota, and Pseudomonadota as the densest phyla in c_AMPs. As a reference, the median of ⍴*_AMP_* for the presented genera is indicated by a magenta dashed line (see ***Tables SI7*** and ***SI8***).

One example of this strain-specific behavior is AMP10.018_194, the only c_AMP found in *Mycoplasma pneumoniae* genomes. *M. pneumoniae* strains are traditionally classified into two groups based on their P1 adhesin gene^53^. Of the 76 *M. pneumoniae* genomes present in our study, 29 were of type-1, 29 were of type-2, and the remaining 18 were of the undetermined type in this classification system^54^ (***Methods - Determination of accessory AMPs***). Twenty-six of the 29 type-2 genomes contained AMP10.018_194, as did 2 undetermined type genomes, but none of the type-1 genomes contained this AMP.

### Bacterial strains from the human gut have more c_AMP genes than conspecific strains from other human body sites

We investigated the taxonomic composition of the AMPSphere by annotating contigs with the GTDB taxonomy^55,56^ (see ***Methods - Differences in the c_AMP density in microbial species from different habitats***), which resulted in 570,187 c_AMPs being annotated to a genus or species. The genera contributing the most c_AMPs to AMPSphere were *Prevotella* (18,593 c_AMPs), *Bradyrhizobium* (11,846 c_AMPs), *Pelagibacter* (6,675 c_AMPs), *Faecalibacterium* (5,917 c_AMPs), and CAG-110 (5,254 c_AMPs) (see ***Fig. 4***). This distribution reflects the fact that these genera are among those that contribute the most assembled sequences in our dataset (all occupying percentiles above 99.75% among the assembled genera; see ***Table SI7***). Therefore, we computed the c_AMP density (⍴*_AMP_*) as the number of c_AMP genes found per megabase pair of assembled sequence. The densest genera were environmental microorganisms, such as *Algorimicrobium* 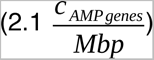, as well as non-cultured taxa, *e.g.*, TMED78 (1.6), SFJ001 (1.5), STGJ01 (1.4), and CAG-462 (1.4). However, when we considered phylum-level annotations, we observed none of the above-mentioned genera belonging to the Methylomirabilota phylum, the group with the highest ⍴*_AMP_*, followed by Fusobacteriota. The absence of these genera is due to the presence of contigs assigned to these phyla, but not to a specific genus, likely because of a lack of representation in the database.

These environmental high c_AMP density genera are, however, low abundance taxa (see ***Methods - Differences in the c_AMP density in microbial species from different habitats***) and, the average *ρ_AMP_* of animal-host-associated samples is 1.6-fold higher than in samples from non-host-associated habitats (P_Mann_ < 10^−330^; ***Fig. 5A*** and ***Fig. SI4***). Differences between habitats reflect both changes in high-level taxonomic composition and subspecies variation. To disentangle these two effects, we analyzed the 3,930 microbial species that were present in at least 10 samples from two different habitats, comparing the AMP density in sequences from the same species in different habitats. This resulted in 1,531 species showing significant differences in at least one pair of habitats (FDR < 0.05, Holm-Sidak; see ***Methods - Differences in the c_AMP density in microbial species from different habitats***). For example, *Prevotella copri* has a higher density when found in the cat and human gut compared to other mammalian hosts or in wastewater (P_Kruskal_ = 4.9·10^−276^, ***Fig. 5B*** and ***Table SI8***). When comparing the human gut and oral cavity (the two body sites with the greatest overlap in species), in general, microbial strains have higher AMP density in the human gut (P_Mann_ = 1.2·10^−4^, N_Species_ = 37, ***Fig. 5C*** and ***Table SI8***). We also checked non-animal hosts, such as plants, and observed that microbial species simultaneously observed in soil and plant-associated microbiome usually have higher AMP density in soil (P_Mann_ = 2.8·10^−4^, N_Species_ = 130, ***Fig. 5D*** and ***Table SI8***).

**Figure 5.**
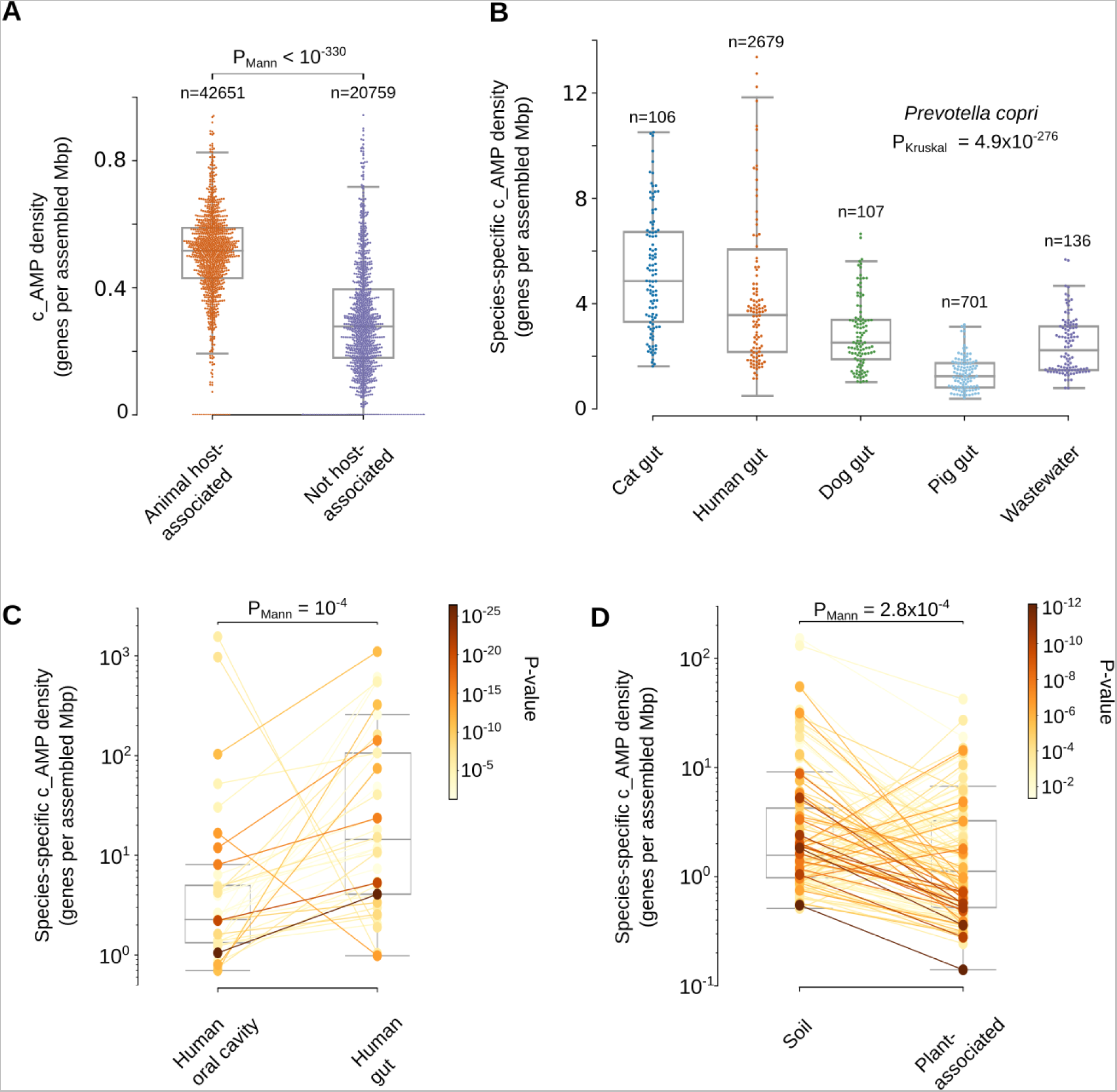
Habitats differ in their c_AMP densities, with differences being observed between conspecific strains. (***A***) Host-associated samples presented a higher ⍴*_AMP_* (calculated at a genus and samples levels) than samples from environmental samples (a random sample of 1 thousand dots for each group was drawn excluding outliers – see ***Differences in the c_AMP density in microbial species from different habitats*** in ***Methods***). (***B***) *Prevotella copri* has a higher ⍴AMP in cat and human guts compared to the same species in the guts of pigs and dogs. 106 randomly selected points are shown for each host. (***C***) Investigating the species-specific ⍴*_AMP_* of microbes found in samples from the human gut and human oral cavity, we observed 34 out of the 37 tested species presenting a higher c_AMP density in the gut. (***D***) The effect of a host when it is not an animal was investigated by verifying the species-specific ⍴*_AMP_* of microbes happening in samples from soil and plants. We observed 85 out of the 130 tested species presenting a higher density in soils. For panels ***C*** and ***D*** the significance was color-encoded using a Log_10_(P_Mann_) scale. See also ***Fig. SI4***, ***Fig. SI5,*** and ***Table SI8***. Our results also showed that differences in ⍴*_AMP_* observed also were kept even when restricting the c_AMP genes by controlling their quality (***Fig. SI5***).

### More transmissible species have lower AMP density

To further establish the importance of AMP production in ecological processes, we investigated the role of AMPs in the mother-to-child transmissibility of bacterial species in a recently published dataset^57^, by correlating the *ρ_AMP_* for each bacterial species to the published measures of microbial transmission. Human gut bacteria showed increased transmissibility at lower AMP densities (R_Spearman_ = −0.44, P_Holm-Sidak_ = 9.5·10^−3^, N_Species_ = 45). Similarly, in human oral microbiome bacterial species, transmissibility from mother to offspring is consistently inversely correlated with their *ρ_AMP_* for the first year (R_Spearman_ = −0.53, P_Holm-Sidak_ = 1.2·10^−3^, N_Species_ = 45), 3 years (R_Spearman_ = −0.38, P_Holm-Sidak_ = 3.9·10^−3^, N_Species_ = 77), and up to 18 years (R_Spearman_ = −0.35, P_Holm-Sidak_ = 3.9·10^−3^, N_Species_ = 91). For the human oral datasets, we observed that lower *ρ_AMP_* correlated to species transmission in the case of people cohabitating (R_Spearman_ = −0.33, P_Holm-Sidak_ = 6.3·10^−3^, N_Species_ = 91). Thus, the AMP contents may influence microbial transmission success rates.

### Physicochemical features and secondary structure of AMPs

To investigate the properties and structure of the synthesized peptides, we first compared their amino acid composition to AMPs from available databases (DRAMP 3.0^44^, DBAASP^58^, and APD3^59^). Overall, the composition was similar, as was expected, given that Macrel’s ML model was trained using known AMPs^40^. Notably, the AMPSphere sequences displayed a slightly higher abundance of aliphatic amino acid residues, specifically alanine and valine. However, these AMPSphere sequences consistently differed (***Fig. 6A***) from encrypted peptides (EPs), which are peptides previously identified within proteins, including those found in the human proteome^3^. The resemblances in amino acid composition between the identified c_AMPs and known AMPs suggested similar physicochemical characteristics and secondary structures, both of which are recognized for their influence on antimicrobial activity^60^. The c_AMPs exhibited comparable hydrophobicity, net charge, and amphiphilicity to AMPs sourced from databases (***Fig. SI1***). Furthermore, they displayed a slight propensity for disordered conformations (***Fig. 6B***) and had a lower positive charge compared to EPs (***Fig. 6A***).

**Figure 6.**
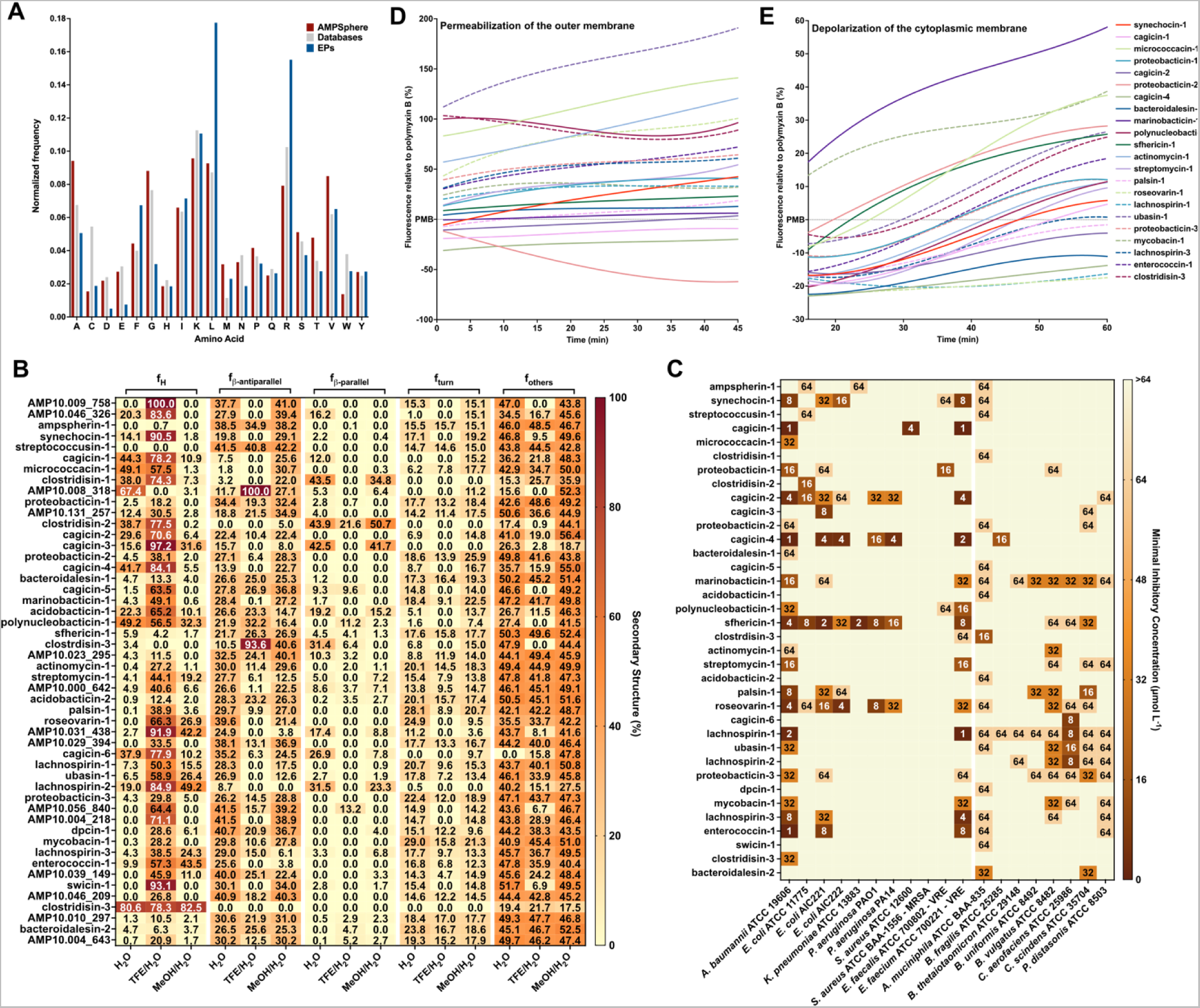
Amino acid composition, structure, antimicrobial activity, and mechanism of action of c_AMPs. ***(A)*** Amino acid frequency in c_AMPs from AMPSphere, AMPs from databases (DRAMP v3, APD3, and DBAASP), and encrypted peptides (EPs) from the human proteome. ***(B)*** Heat map with the percentage of secondary structure found for each peptide in three different solvents: water, 60% trifluoroethanol in water, and 50% methanol in water. Secondary structure was calculated using BeStSel server^130^. ***(AC)*** Activity of c_AMPs assessed against ESKAPEE pathogens and human gut commensal strains. Briefly, 10^6^ CFU·mL^−1^ was exposed to c_AMPs two-fold serially diluted ranging from 64 to 1 μmol·L^−1^ in 96-wells plates and incubated at 37 °C for one day. After the exposure period, the absorbance of each well was measured at 600 nm. Untreated solutions were used as controls and minimal concentration values for complete inhibition were presented as a heat map of antimicrobial activities (μmol·L^−1^) against 11 pathogenic and eight human gut commensal bacterial strains. All the assays were performed in three independent replicates and the heatmap shows the mode obtained within the two-fold dilutions concentration range studied. ***(D)*** Fluorescence values relative to polymyxin B (PMB, positive control) of the fluorescent probe 1-(N-phenylamino)naphthalene (NPN) that indicate outer membrane permeabilization of *A. baumannii* ATCC 19606 cells. ***(E)*** Fluorescence values relative to PMB (positive control) of 3,3′-dipropylthiadicarbocyanine iodide [DiSC3-(5)], a hydrophobic fluorescent probe, used to indicate cytoplasmic membrane depolarization of *A. baumannii* ATCC 19606 cells. Depolarization of the cytoplasmic membrane occurred with a slow kinetics compared to the permeabilization of the outer membrane and took approximately 20 min to stabilize.

Subsequently, we conducted experimental assessments of the secondary structure of the active c_AMPs using circular dichroism (***Fig. 6B*** and ***SI6***). Similar to AMPs documented in databases, peptides derived from AMPSphere exhibited a pronounced propensity for adopting α-helical structures. Notably, they also displayed an unusually high content of β-antiparallel structure in both water and methanol/water mixtures (***Fig. 6B***), despite their amino acid composition similarities to AMPs and EPs. We attribute these findings to the slightly elevated occurrence of alanine and valine residues, which are known to favor β-like structures with a preference for β-antiparallel conformation^61^.

### Validation of c_AMPs as potent AMPs through in vitro assays

To evaluate the potential antimicrobial properties of c_AMPs, we selected and chemically synthesized 50 peptide sequences based on their abundance, predicted solubility, and taxonomic diversity (***Methods*** - ***Selection of peptides to synthesis and activity testing***). Next, we subjected these peptides to testing against 11 clinically relevant pathogenic strains, encompassing *Acinetobacter baumannii*, *Escherichia coli* (including one colistin-resistant strain), *Klebsiella pneumoniae*, *Pseudomonas aeruginosa*, *Staphylococcus aureus* (including one methicillin-resistant strain), vancomycin-resistant *Enterococcus faecalis*, and vancomycin-resistant *Enterococcus faecium*. Our initial screening revealed that 27 AMPs (54% of the total synthesized) completely eradicated the growth of at least one of the pathogens tested (***Fig. 6C***). Remarkably, in some cases, the AMPs were active at concentrations as low as 1 μmol·L^−1^. Several of the Gram-negative bacteria, *i.e.*, *A. baumannii, E. coli,* and *P. aeruginosa*, as well as the Gram-positive strain of vancomycin-resistant *E. faecium*, displayed higher susceptibility to the AMPs, with 22, 15, 4 and 15 peptide hits, respectively. However, none of the tested AMPs targeted methicillin-resistant *S. aureus* (MRSA) (***Fig. 6C***).

### The growth of human gut commensals is impaired by c_AMPs

We screened the AMPs against eight of the most prevalent members of the human gut microbiota. We tested commensal bacteria belonging to four phyla (Verrucomicrobiota, Bacteroidota, Actinomycetota, and Bacillota), *i.e*., *Akkermansia muciniphila*, *Bacteroides fragilis*, *Bacteroides thetaiotaomicron*, *Bacteroides uniformis*, *Phocaeicola vulgatus (formerly Bacteroides vulgatus)*, *Collinsella aerofaciens*, *Clostridium scindens*, and *Parabacteroides distasonis*.

While it is commonly observed that known natural AMPs do not target microbiome strains^62^, our study found that 30 of the synthesized AMPs (60%) demonstrated inhibitory effects on at least one commensal strain at low concentrations levels (8-16 μmol·L^−1^). Although this concentration range was higher than that required to inhibit pathogens (1-4 μmol·L^−1^), it still falls within the highly active range of AMPs based on previous studies^63–65^ (***Fig. 6C***). Interestingly, all the analyzed gut microbiome strains were susceptible to at least two c_AMPs, with strains of *A. muciniphila*, *B. uniformis*, *P. vulgatus*, *C. aerofaciens*, *C. scindens*, and *P. distasonis* exhibiting the highest susceptibility. In total, 36 AMPs (72% of the total synthesized peptides) demonstrated antimicrobial activity against pathogens and/or commensals.

### Depolarization and permeabilization of the bacterial membrane by AMPs from AMPSphere

To gain insights into the mechanism of action responsible for the antimicrobial activity observed in the peptides derived from AMPSphere (***Fig. 6C***), we conducted experiments to assess their ability to permeabilize and depolarize the cytoplasmic and outer membranes of bacteria at their Minimum Inhibitory Concentrations - MICs. Specifically, we investigated the effects of 22 peptides on *A. baumannii* (***Fig. 6D-E*** and ***Fig. SI7A,D***) and 4 peptides on *P. aeruginosa* (***Fig. SI7B-C,E***). For comparison, we used polymyxin B, a peptide antibiotic known for its membrane permeabilization and depolarization properties, as a control in these experiments^3^.

To investigate the potential permeabilization of the outer membranes of Gram-negative bacteria by active AMPs derived from AMPSphere, we conducted 1-(N-phenylamino)naphthalene (NPN) uptake assays (***Fig. SI7A-B***). NPN is a lipophilic fluorophore that exhibits increased fluorescence in the presence of lipids found in bacterial outer membranes. The uptake of NPN indicates membrane permeabilization and damage. Among the 22 peptides evaluated for activity against *A. baumannii*, 10 peptides caused significant permeabilization of the outer membrane, resulting in fluorescence levels at least 50% higher than that of polymyxin B (***Fig. 6D***). Only three peptides exhibited lower permeabilization than polymyxin B (***Fig. 6D***). In the case of *P. aeruginosa* cells, two out of the four tested peptides showed higher permeabilization than polymyxin B (***Fig. SI7C***).

To evaluate the potential membrane depolarization effect of the AMPs from AMPSphere, we utilized the fluorescent dye 3,3′-dipropylthiadicarbocyanine iodide [DiSC_3_-(5)] (***Fig. SI7D-E***). Among the peptides tested against *A. baumannii* and *P. aeruginosa*, marinobacticin-1 (AMP10.321_460) and cagicin-2 (AMP10.014_861) exhibited greater cytoplasmic membrane depolarization than polymyxin B (***Fig. 6E***;***Fig. SI7E-F***), respectively. Interestingly, all the tested AMPSphere peptides displayed a characteristic crescent-shaped depolarization pattern compared to polymyxin B, with lower levels of depolarization observed during the first 20 minutes of exposure, followed by an increase in depolarization over time (***Fig. 6E*** and ***Fig. 7D-F***). Taken together, these results indicate that the kinetics of cytoplasmic membrane depolarization are slower compared to the kinetics of outer membrane permeabilization, which occurs rapidly upon interaction with the bacterial cells.

**Figure 7.**
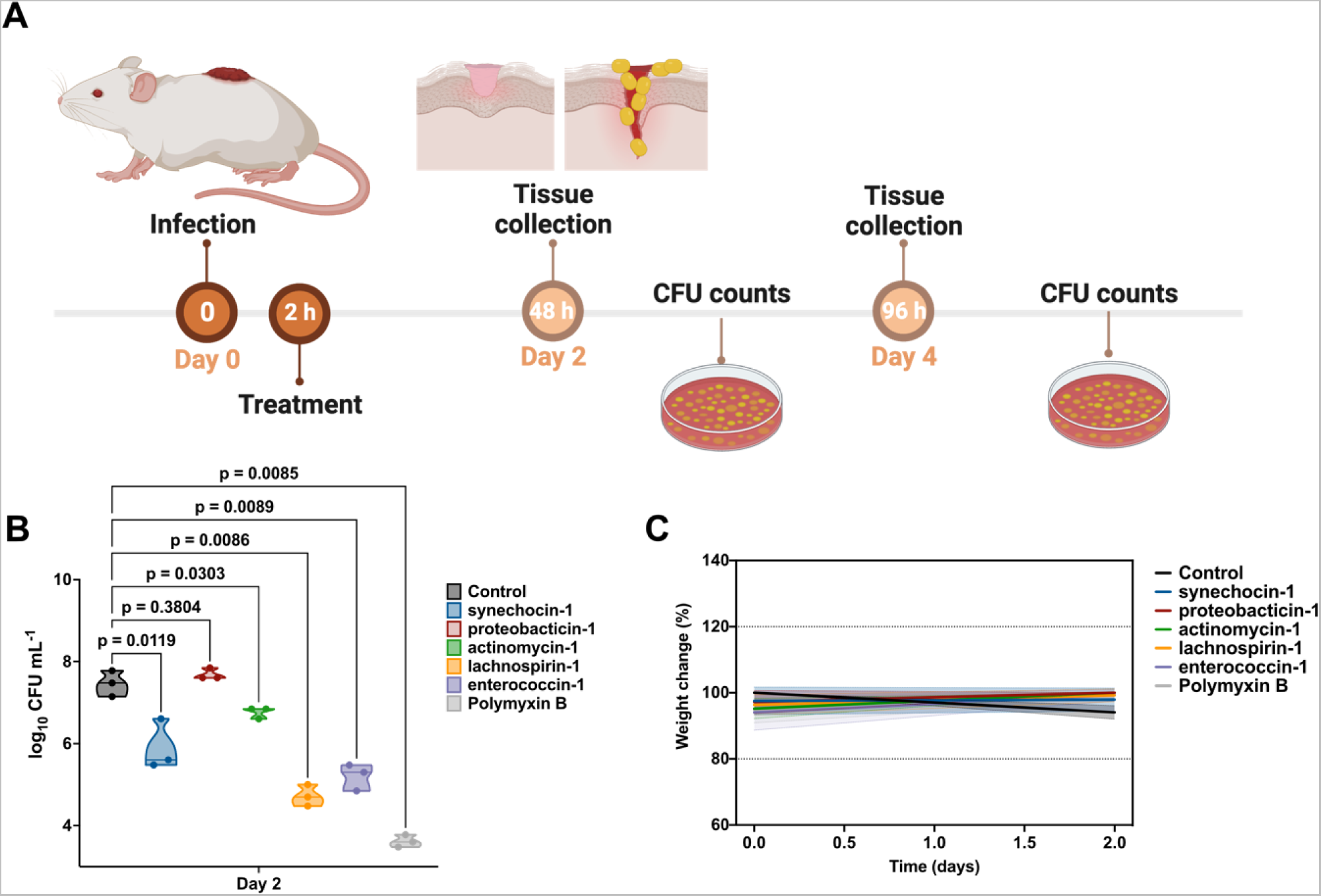
Anti-infective activity of AMPs in pre-clinical animal model. ***(A)*** Schematic of the skin abscess mouse model used to assess the anti-infective activity of the peptides against *A. baumannii* cells. ***(B)*** Peptides were tested at their MIC in a single dose one hour after the establishment of the infection. Each group consisted of three mice (n = 3) and the bacterial loads used to infect each mouse derived from a different inoculum. ***(C)*** To rule out toxic effects of the peptides, mouse weight was monitored throughout the experiment. Statistical significance in ***(B)*** was determined using one-way ANOVA where all groups were compared to the untreated control group; P-values are shown for each of the groups. Features on the violin plots represent median and upper and lower quartiles. Data in ***(C)*** are the mean ± the standard deviation. Figure created in BioRender.com.

Our findings indicate that the AMPs from AMPSphere primarily exert their effects by permeabilizing the outer membrane rather than depolarizing the cytoplasmic membrane, revealing a similar mechanism of action to that observed for classical AMPs and EPs.

### AMPs exhibit anti-infective efficacy in a mouse model

Next, we tested the anti-infective efficacy of AMPSphere-derived peptides in a skin abscess murine infection model (***Fig. 7A***). Mice were subjected to infection with *A. baumannii*, a Gram-negative pathogen known for causing severe infections in various body sites including bloodstream, lungs, urinary tract, and wounds. This pathogen is recognized as one of the most dangerous^66^. Five lead AMPs from different sources displayed potent activity against *A. baumannii*: synechocin-1 (AMP10.000_211, 8 μmol·L^−1^) from *Synechococcus* sp. (coral associated, marine microbiome), proteobacticin-1 (AMP10.048_551, 16 μmol·L^−1^) from Pseudomonadota (plant and soil microbiome), actynomycin-1 (AMP10.199_072, 64 μmol·L^−1^) from *Actinomyces* (human mouth and saliva microbiome), lachnospirin-1 (AMP10.015_742, 2 μmol·L^−1^) from *Lachnospira* sp. (human gut microbiome), and enterococcin-1 (AMP10.051_911, 1 μmol·L^−1^) from *Enterococcus faecalis* (human gut microbiome).

The skin abscess infection was established with a bacterial load of 20 μL of *A. baumannii* cells at 1β10^6^ CFU·mL^−1^ onto the wounded area of the dorsal epidermis (***Fig. 7A***). A single dose of each peptide, at their respective MIC (***Fig. 6A***), was administered to the infected area. Two days post-infection, synechocin-1 and actynomycin-1 peptides presented bacteriostatic activity, inhibiting the proliferation of *A. baumannii* cells, whereas lachnospirin-1 and enterococcin-1 presented bactericidal activity similar to that of the antibiotic polymyxin B (MIC = 0.25 μmol·L^−1^), reducing the colony-forming units (CFU) counts up to 3-4 orders of magnitude (***Fig. 7B***). Four days post-infection, none of the AMPs nor polymyxin B achieved a statistically significant reduction of *A. baumannii* growth at the infection site, although treatment with proteobactin-1, lachnospiracin-1, enterococcin-1, and polymyxin B reduced the CFU counts by 1-2 orders of magnitude compared to the untreated control. These results are promising since the AMPSphere peptides were administered only once immediately after the establishment of the abscess, highlighting their anti-infective potential.

Mouse weight was monitored as a proxy for toxicity and no significant changes were observed (***Fig. 7C***), suggesting that the peptides tested are not toxic.

## Discussion

Here, we use ML to identify thousands of novel candidate AMPs in the global microbiome. Building on previous studies that focused specifically on the human gut microbiome^5,36,67^, we cataloged AMPs from the global microbiome across 63,410 publicly available metagenomes, as well as 87,920 high-quality microbial genomes from the ProGenomes v2 database^41^, leading to the creation of AMPSphere (https://ampsphere.big-data-biology.org/), a publicly available resource encompassing 863,498 non-redundant peptides and 6,499 high-quality AMP families from 72 different habitats, including marine and soil environments and the human gut. We show that most of the c_AMPs (91.5%) were previously unknown, lacking detectable homologs in other databases, and about one infive could be detected in independent sets of meta-transcriptomes or metaproteomes.

Two evolutionary mechanisms by which AMPs may be generated were explored. First, mutations in genes encoding longer proteins could generate gene fragments. Among the enriched ortholog groups of proteins from GMGCv1^46^ homologous to c_AMPs, we observed that a majority of groups had unknown function (53.8%), similar to what was reported by Sberro et al.^36^ for small proteins from the human gut microbiome. The second mechanism is that a gene duplication could be followed by mutation, which we observed in the case of ribosomal proteins. Ribosomal proteins can harbor antimicrobial activity^50^, possibly due to their amyloidogenic properties^68^. Nonetheless, the majority of identified AMPs do not have detectable homology to other sequences, highlighting their novelty. The lack of observed homology, however, may be due to limitations in our ability to robustly detect these homology relationships in small sequences, but there is also the possibility that small proteins, such as AMPs, may be more likely to be generated *de novo* and may have repeatedly evolved in various taxa^69^.

Four out of the five genera with the most c_AMPs present in AMPSphere share a host-associated lifestyle. Three of these (*Prevotella*, *Faecalibacterium*, and *CAG-110*) are common in animal hosts (***Fig. 4***). The greater density of c_AMP genes in genera from the Bacillota and Bacillota A phyla is consistent with the well-known diversity of ABC transporters dedicated to the translocation of AMPs found in that group, resulting in improved resistance to compounds that bind extracellular targets; for a review, see Gebhard^70^.

We observed that c_AMPs from AMPSphere are habitat-specific and mostly accessory members of microbial pangenomes. Moreover, species-specific density (⍴*_AMP_*) shows that the habitat plays an important role in shaping c_AMP content. The ⍴*_AMP_* of strains from the same species can differ even across body sites. In particular, we observed higher ⍴*_AMP_* in the human gut compared to the human oral cavity, in agreement with a recent report of a strain-specific AMP (cutymicin), which is present in only some of the hair follicles in the same human host^71^.

Valles-Colomer et al.^57^, who recently analyzed a large collection of human-associated metagenomes, provide a species-specific index of transmissibility for the several transmission scenarios they study (*e.g*., mother to infant). Hypothesizing that AMP production may be related to transmission, we correlated the species-specific ⍴*_AMP_* calculated in AMPSphere with transmission scores. In both the human gut and oral microbiomes, species with higher ⍴*_AMP_* are less transmissible, possibly because AMPs confer protection against strain replacement. Taken together, our results and those of Valles-Colomer et al.^57^ validate the AMP density concept and the applicability of AMPSphere resources to study mechanisms of microbial establishment and competition.

Finally, we experimentally validated predictions made by our ML model and found that 72% (32 out of the 50) synthesized AMPs displayed antimicrobial activity against either pathogens or commensals. Notably, four peptides (cagicin-1, cagicin-4, and enterococcin-1 against *A. baumannii*; and cagicin-1 and lachnospirin-1 against vancomycin-resistant *E. faecium*) presented MIC values as low as 1 μmol·L^−1^, comparable to the MICs of some of the most potent peptides previously described in the literature^64,65^.

We show that AMPs from the AMPSphere tended to target clinically relevant Gram-negative pathogens and also showed activity against vancomycin-resistant *E. faecium*. Although conventional AMPs do not target microbiome bacteria^62^, AMPs from AMPSphere showed efficacy against these bacteria, suggesting potential ecological implications of peptides as protective agents for their producing organisms. Notably, the amino acid composition and physicochemical characteristics of the c_AMPs in AMPSphere differed from those recently identified in EPs^3^.

Moreover, three peptides exhibited anti-infective efficacy in a murine infection model, with lachnospirin-1 and enterococcin-1 being the most potent, resulting in a reduction of bacterial load by up to four orders of magnitude. The active peptides included those derived from both human-associated and environmental microbiota, validating our approach of investigating the global microbiome. Overall, our findings unveil a wide array of novel AMP sequences, highlighting the potential of machine learning in the identification of much-needed antimicrobials.

## Acknowledgments

We thank Marija Dmitrijeva (University of Zurich) for her helpful comments on a previous version of the manuscript. We thank members of the Coelho group and de la Fuente Lab for insightful discussions. Cesar de la Fuente-Nunez holds a Presidential Professorship at the University of Pennsylvania and acknowledges funding from the Procter & Gamble Company, United Therapeutics, a BBRF Young Investigator Grant, the Nemirovsky Prize, Penn Health-Tech Accelerator Award, and the Dean’s Innovation Fund from the Perelman School of Medicine at the University of Pennsylvania. We thank Dr. Mark Goulian for kindly donating the following strains: *Escherichia coli* AIC221 [*Escherichia coli* MG1655 phnE_2::FRT (control strain for AIC 222)] and *Escherichia coli* AIC222 [*Escherichia coli* MG1655 pmrA53 phnE_2::FRT (polymyxin resistant)]. This work was funded by the following grants: National Key R&D Program of China (2020YFA0712403, 2018YFC0910500) (LPC, ZXM); National Natural Science Foundation of China (61932008, 61772368) (LPC, ZXM); Shanghai Science and Technology Innovation Fund (19511101404) (LPC, ZXM); Shanghai Municipal Science and Technology Major Project (2018SHZDZX01) (LPC, ZXM); The Science and Technology Commission of Shanghai Municipality (22JC1410900) (LPC); Langer Prize (AIChE Foundation) (CFN); National Institutes of Health grant R35GM138201 (CFN); Defense Threat Reduction Agency grant HDTRA11810041 (CFN); Defense Threat Reduction Agency grant HDTRA1-21-1-0014 (CFN); Defense Threat Reduction Agency grant HDTRA1-23-1-0001 (CFN); PID2021-127210NB-I00, MCIN/AEI/10.13039/501100011033/FEDER, UE. (JHC); “La Caixa” Foundation (ID 100010434), fellowship code LCF/BQ/DI18/11660009 (ARdR); European Union’s Horizon 2020 research and innovation program under the Marie Skłodowska-Curie grant agreement 713673 (ARdR).

## Declaration of interests

Cesar de la Fuente-Nunez provides consulting services to Invaio Sciences and is a member of the Scientific Advisory Boards of Nowture S.L. and Phare Bio. The de la Fuente Lab has received research funding or in-kind donations from United Therapeutics, Strata Manufacturing PJSC, and Procter & Gamble, none of which were used in support of this work. All other authors state they do not have any competing interests.

## Author contributions

1. ***Conceptualization:*** CDSJ LPC MDTT CFN
2. ***Data curation:*** CDSJ YD TSBS AF LPC MDTT CFN
3. ***Formal analysis:*** CDSJ LPC MDTT
4. ***Funding acquisition:*** LPC XMZ CFN
5. ***Investigation:*** CDSJ LPC MDTT CFN
6. ***Methodology:*** CDSJ YD JHC ARdR LPC MDTT CFN
7. ***Project administration:*** LPC XMZ PB CFN
8. ***Resources:*** LPC XMZ CFN
9. ***Supervision:*** LPC CFN
10. ***Visualization:*** CDSJ JHC JS AV AH CZ LPC MDTT
11. ***Writing – original draft:*** CDSJ LPC
12. ***Writing – review & editing:*** CDSJ YD JHC ARdR TSBS AF PB XMZ LPC MDTT CFN

## KEY RESOURCES TABLE

| **REAGENT or RESOURCE** | **SOURCE** | **IDENTIFIER** |
| --- | --- | --- |
| **Bacterial and virus strains** |  |  |
| *Acinetobacter baumannii* | American Type Culture Collection | ATCC 19606 |
| *Escherichia coli* | American Type Culture Collection | ATCC 11775 |
| *Escherichia coli* | *Escherichia coli*<br>MG1655<br>phnE_2::FRT | AIC221 |
| *Escherichia coli* | *Escherichia coli* | AIC222 |
|  | MG1655 pmrA53<br>phnE_2::FRT<br>(polymyxin-resistant;<br>colistin-resistant<br>strain) |  |
| *Klebsiella pneumoniae* | American Type<br>Culture Collection | ATCC 13883 |
| *Pseudomonas aeruginosa* |  | PA01 |
| *Pseudomonas aeruginosa* |  | PA14 |
| *Staphylococcus aureus* | American Type<br>Culture Collection | ATCC 12600 |
| *Staphylococcus aureus* | American Type<br>Culture Collection | ATCC BAA-1556<br>(methicillin-resistant strain) |
| *Akkermansia muciniphila* | American Type<br>Culture Collection | ATCC BAA-635 |
| *Bacteriodes fragilis* | American Type<br>Culture Collection | ATCC 25285 |
| *Bacteroides thetaiotaomicron* | American Type<br>Culture Collection | ATCC 29148 |
| *Bacteroides uniformis* | American Type<br>Culture Collection | ATCC 8492 |
| *Bacteroides vulgatus (Phocaeicola vulgatus)* | American Type<br>Culture Collection | ATCC 8482 |
| *Collinsella aerofaciens* | American Type<br>Culture Collection | ATCC 25986 |
| *Clostridium scindens* | American Type<br>Culture Collection | ATCC 35704 |
| *Parabacteroides distasonis* | American Type | ATCC 8503 |
|  | Culture Collection |  |
| ***Chemicals*** |  |  |
| Luria-Bertani broth | BD | 244620 |
| Tryptic soy broth | Sigma | T8907-1KG |
| Agar | Sigma | 05039 |
| MacConkey agar | RPI | M42560-500.0 |
| Phosphate buffer saline | Sigma | P3913-10PAK |
| Glucose | Sigma | G5767 |
| 1-(N-phenylamino)naphthalene | Sigma | 104043 |
| 3,3'-dipropylthiadicarbocyanine iodide | Sigma | 43608 |
| HEPES | Fisher | BP310-100 |
| Potassium chloride (KCl) | Sigma | P3911 |
| ***Deposited data*** |  |  |
| Code for generation of AMPSphere | This study | https://github.com/BigDataBiology/AMPSphere_manuscript |
| AMPSphere database | This study | https://zenodo.org/record/4606582 |
| ***Software and algorithms*** |  |  |
| NGLess 1.3.0 | Coelho et al.<sup>72</sup> | https://github.com/ngless- |
|  |  | toolkit/ngless |
| JUG 2.1.1 | Coelho<sup>73</sup> | https://github.com/luispedro/jug |
| Prodigal 2.6.3 | Hyatt et al.<sup>32</sup> | https://github.com/hyattpd/Prodigal |
| Macrel v.1.0.0 | Santos-Júnior et al.<sup>40</sup> | https://github.com/BigDataBiology/macrel |
| CDHit 4.8.1 | Fu et al.<sup>74</sup> | https://github.com/weizhongli/cdhit |
| MMseqs2 | Steinegger and Söding<sup>75</sup> | https://github.com/soedinglab/MMseqs2 |
| python 3.8.2 | Van Rossum<sup>76</sup> | https://www.python.org/ |
| matplotlib 3.4.3 | Hunter<sup>77</sup> | https://matplotlib.org/ |
| numpy 1.21.2 | Harris et al.<sup>78</sup> | https://numpy.org/ |
| pandas 1.3.2 | McKinney<sup>79</sup> | https://pandas.pydata.org/ |
| plotly 5.2.1 | Plotly Technologies Inc, 2015 | https://plot.ly |
| scipy 1.7.1 | Virtanen et al.<sup>80</sup> | https://www.scipy.org |
| scikit-learn 0.24 | Pedregosa et al.<sup>81</sup> | https://scikit- |
|  |  | learn.org/ |
| scikit-bio 0.5.6 | The scikit-bio development team, 2020<sup>82</sup> | http://scikit-bio.org/ |
| BioPython 1.7.9 | Cock et al.<sup>83</sup> | https://biopython.org/ |
| eggnog-mapper v2 | Cantalapiedra et al.<sup>84</sup> | https://github.com/eggnogdb/eggnog-mapper |
| HMMer 3.3+dfsg2-1 | Eddy<sup>85</sup> | http://hmmer.org/ |
| modIAMP 4.2.2 | Müller et al.<sup>86</sup> | https://pypi.org/project/modlamp/ |
| FastTree 2.1 | Price et al.<sup>87</sup> | www.microbesonline.org/fasttree/ |
| FastANI v.1.33 | Jain et al.<sup>88</sup> | https://github.com/ParBLiSS/FastANI |
| Megahit 1.2.9 | Li et al.<sup>89</sup> | https://github.com/voutcn/megahit/ |
| **Other** |  |  |
| ProGenomes2 | Mende et al.<sup>41</sup> | http://progenomes.embl.de/ |
| DRAMP - Data repository of antimicrobial peptides 3.0 | Shi et al.<sup>44</sup> | http://dramp.cpu-bioinfor.org/ |
| UniprotKB 2021_03 | The UniProt Consortium<sup>90</sup> | https://www.uniprot.org/ |
| Eggnog v.5.0 | Huerta-Cepas et al.<sup>48</sup> | http://eggnog5.embl.de/ |
| SmProt database v.2.0 | Hao et al.<sup>45</sup> | http://bigdata.ibp.ac.cn/SmProt/index.html |
| StarPep45k | Aguilera-Mendoza et al.<sup>91</sup> | http://mobiosd-hub.com/starpep |
| PFAM 34.0 | Mistry et al.<sup>92</sup> | http://pfam.xfam.org/ |
| AntiFAM v.6.0 | Eberhardt et al.<sup>93</sup> | https://www.ebi.ac.uk/research/bateman/software/antifam-tool-identify-spurious-proteins |
| GTDB 07-RS95 | Parks et al.<sup>55,56</sup> | https://gtdb.ecogenomic.org/ |
| NCBI release 207 | NCBI Resource Coordinators<sup>94</sup> | https://ftp.ncbi.nih.gov/refseq/release/ |
| DBAASP | Pirtskhalava et al.<sup>58</sup> | https://dbaasp.org/home |
| APD3 | Wang and Wang<sup>59</sup> | https://aps.unmc.edu/ |

## RESOURCE AVAILABILITY

### Lead contact

Further information and requests for resources and reagents should be directed to and will be fulfilled by the lead contacts: Cesar de la Fuente-Nunez and Luis Pedro Coelho.

### Materials availability

Peptides were obtained from AAPPTec and synthesized using solid-phase peptide synthesis and 9-fluorenylmethoxycarbonyl (Fmoc) strategy.

### Data and code availability

- Metagenomes and Genomes data are publicly available at the European Nucleotide Archives (ENA) as of the date of publication. Accession numbers are listed in the supplementary tables.
- All original code has been deposited at GitHub in the link: github.com/BigDataBiology/AMPsphere_manuscript
- AMPSphere files have been deposited in Zenodo and are publicly available as of the date of publication. DOIs are listed in the key resources table.
- AMPSphere is also available as a public online resource: https://ampsphere.big-data-biology.org/
- Any additional information required to reanalyze the data reported in this paper is available from the lead contact upon request.

## EXPERIMENTAL MODEL AND SUBJECT DETAILS

### Bacterial strains and growth conditions

The pathogenic strains *Acinetobacter baumannii* ATCC 19606, *Escherichia coli* ATCC 11775, *Escherichia coli* AIC221 [*Escherichia coli* MG1655 phnE_2::FRT (control strain for AIC 222)], *Escherichia coli* AIC222 [*Escherichia coli* MG1655 pmrA53 phnE_2::FRT (polymyxin-resistant; colistin-resistant strain)], *Klebsiella pneumoniae* ATCC 13883, *Pseudomonas aeruginosa* PA01, *Pseudomonas aeruginosa* PA14, *Staphylococcus aureus* ATCC 12600, *Staphylococcus aureus* ATCC BAA-1556 (methicillin-resistant strain), *Enterococcus faecalis* ATCC 700802 (vancomycin-resistant strain), and *Enterococcus faecium* ATCC 700221 (vancomycin-resistant strain) were grown and plated on Luria-Bertani (LB) agar plates and incubated overnight at 37°C from frozen stocks. After incubation, one isolated colony was transferred to 6 mL of medium (LB), and cultures were incubated overnight (16 h) at 37°C. The following day, inocula were prepared by diluting the overnight cultures 1:100 in 6 mL of the respective media and incubating them at 37°C until bacteria reached logarithmic phase (OD_600_ = 0.3-0.5).

The gut commensal strains *Akkermansia muciniphila* ATCC BAA-635, *Bacteroides fragilis* ATCC 25285, *Bacteroides thetaiotaomicron* ATCC 29148, *Bacteroides uniformis* ATCC 8492, *Bacteroides vulgatus* ATCC 8482 (*Phocaeicola vulgatus*), *Collinsella aerofaciens* ATCC 25986, *Clostridium scindens* ATCC 35704, and *Parabacteroides distasonis* ATCC 8503 were grown in brain heart infusion (BHI) agar plates enriched with 0.1% (v:v) vitamin K3 (1 mg·mL^−1^), 1% (v:v) hemin (1 mg·mL^−1^, diluted with 10 mL of 1 N sodium hydroxide), and 10% (v:v) L-cysteine (0.05 mg·mL^−1^), from frozen stocks and incubated overnight at 37°C. Resazurin was used as oxygen indicator. After the incubation period, a single isolated colony was transferred to 3 mL of BHI broth and incubated overnight at 37°C. The next day, inocula were prepared by diluting the bacterial overnight cultures 1:100 in 3 mL of BHI broth and incubated at 37°C until cells reached the logarithmic phase (OD _600_ = 0.3-0.5).

### Skin abscess infection mouse model

To assess the anti-infective efficacy of the peptides against *A. baumannii* ATCC 19606 in a skin abscess infection mouse model, the bacteria were cultured in tryptic soy broth (TSB) medium until an OD_600_ of 0.5 was reached. Next, the cells were washed twice with sterile PBS (pH 7.4) and suspended to a final concentration of 5·10^6^ colony-forming units (CFU) per mL^−1^. Six-week-old female CD-1 mice, after being anesthetized with isoflurane, were subjected to a superficial linear skin abrasion on their backs in an area that they could not touch with their mouth or limbs. An aliquot of 20 μL containing the bacterial load was then administered over the abraded area. A single dose of the peptides diluted in water at their MIC value was administered to the infected area two h after the infection. The animals were euthanized two-and four-days post-infection, and the infected area was extracted and homogenized for 20 min using a bead beater (25 Hz) and 10-fold serially diluted for CFU quantification on MacConkey agar plates for easy differentiation of *A. baumannii* colonies. The experimental groups consisted of 3 mice per group (n = 3), and each mouse was infected with an inoculum from a different colony to ensure variability.

## METHOD DETAILS

### Selection of metagenomes and high-quality microbial genomes

Selection of metagenomes and genomes to compose the AMPSphere was similar to that adopted by Coelho et al.^46^. Only public metagenomes on 1 January 2020 produced with Illumina instruments (except for MiSeq), with at least 2 million reads and, on average, 75 bp long, were downloaded from the European Nucleotide Archive (ENA). These samples met two criteria: (1) tagged with taxonomy ID 408169 (for metagenome) or is a descendent of it in the taxonomic tree; and/or (2) experiments with the library source listed as “METAGENOMIC”. Samples were grouped by project and all projects with at least 20 samples were considered. Additionally, studies based on a whitelist, including metagenomes deposited by the Integrated Microbial Genomes System (IMG) missing from ENA, also were included. Metadata was manually curated from their describing literature and Biosamples database^95^. The habitat classification took into account the metadata, creating groups based on the similarity of habitat conditions, such as air, anthropogenic, aquatic, host-associated, ph:alkaline, sediment, terrestrial, and others. The sample origins and other relevant information related to host species were assessed using the NCBI taxonomic identification number. High-quality microbial genomes were selected from ProGenomes2 database^41^. The resulting 63,410 publicly available metagenomes and 87,920 high-quality microbial genomes are listed in ***Table SI1***.

### Reads trimming and assembly

Reads were processed using NGLess^72^, trimming positions with quality lower than 25, and discarding reads shorter than 60 bp post-trimming. Metagenomes obtained from a host-associated microbiome passed through a filtering of reads mapping to the host genome when available. Reads were assembled with MEGAHIT 1.2.9^89^ and the taxonomy of 16,969,685,977 contigs assembled from more than 14.7 trillion base pairs of sequenced DNA was inferred as previously described^96^, using MMSeqs2^75^ to map the sequences against the GTDB release 95^55,56^, later manually curated to conform to the International Code of Nomenclature of Prokaryotes.

### smORF and AMP prediction

Analogously to Sberro et al.^36^, we used a modified version of Prodigal^32^ to predict smORFs (33 to 303 bp) from contigs. The 4,599,187,424 redundant smORFs, most of which (99.25%) originated in metagenomes were then de-duplicated, yielding 2,724,621,233 non-redundant smORFs. Macrel^40^ was run on the de-duplicated smORFs to predict c_AMPs. Singleton sequences (those appearing in a single sample or genome) were eliminated, except when they had a significant match (Amino acids identity ≥ 75% and E-value ≤ 10^−5^) to a sequence from the Data Repository of Antimicrobial Peptides - DRAMP 3.0^44^ using the ‘easy-search’ method from MMSeqs2^75^.

AMP genes originating from ProGenomes2^41^ had the taxonomy of the original genome assigned to them, whereas the genes from metagenomes were assigned the taxonomy predicted for the contig where they were found. Insights about potential structural conformations were obtained by using the function secondary_structure_fraction from ProtParam module implemented in the SeqUtils in Biopython^83^. It calculates the fraction of amino acids which tend to assume conformations of helix [VIYFWL], turn [NPGS], and sheet [EMAL].

AMPSphere encompassed a total of 863,498 non-redundant predicted c_AMPs encoded by 5,518,294 redundant genes. AMP densities were estimated as the number of AMPs per assembled base pairs in a sample or a species.

### AMP families

Protein families have a relationship between the structure, function, and origin of proteins^97^. We used a reduced amino acids alphabet of 8 letters^47^ - [LVIMC], [AG], [ST], [FYW], [EDNQ], [KR] - to cluster together sequences which are likely to have similar functions^47,98^. These c_AMPs were hierarchically clustered in this reduced alphabet using three sequential identity cutoffs (100%, 85%, and 75%) with CD-Hit^74^ to improve the profiling and pattern searching at the protein space^99^. A cluster was considered an AMP family when it consisted of at least 8 sequences^36^. Representative sequences of peptide clusters were selected according to their length (taking the longest) with ties being broken by considering alphabetical order.

To validate the clustering procedure, we used a sample of 3,000 thousand sequences randomly drawn from AMPSphere, excluding the representatives. These sequences were aligned against the representative sequence of their cluster using the Smith-Waterman algorithm^100^ with the blocks substitution matrix 62 (BLOSUM 62), and a gap open and extension penalties of −10 and −0.5, respectively. The alignment score was then converted to an E-value, according to the model by Karlin and Altschul^101^, using the values of κ (0.132539) and λ (0.313667) constants adjusted to search for a short input sequence as implemented in the BLAST algorithm^102,103^. Alignments were considered significant when their E-value was less than 10^−5^. We found that more than 95.3% of alignments produced in the first two levels (100% and 100-85% of identity) were significant along with 77.1% of those from the third level (100-85-75% of identity) – see ***Fig. SI3***.

### Quality control of c_AMPs

The genes of c_AMPs were subjected to 5 different quality tests to reduce the likelihood that the observed peptides were artifacts or fragments of larger proteins. Initially, the peptides were searched against AntiFam v.7.0^93^ using HMMSearch^85^ with the option “--cut_ga”, and significant hits were classified as spurious. We observed that 99.9% of the c_AMPs in AMPSphere do not belong to AntiFAMs.

A test for the terminal positioning in contigs checked if there were in-frame stop codons upstream to the smORF coding for a given c_AMP. When no stop codon is found, we cannot rule out the possibility that the smORF is part of a larger gene due to a fragmentary assembly. Most (68.4%) of the c_AMPs are encoded by at least a gene that is not terminally placed.

The RNAcode program^104^ predicts protein-coding regions based on evolutionary signatures typical for protein genes. This analysis depends on a set of homologous and non-identical genes. Therefore, AMP clusters containing at least 3 gene variants were aligned. Given that an extensive portion of the AMPSphere candidates - 53% (459,910 out of 863,498) is not part of such a cluster, they could not be tested. Of the tested c_AMPs, 53% (215,421 out of 403,588) were considered genes with evolutionary traits of protein-coding sequences.

To further verify the experimental evidence of c_AMPs, we checked for evidence of transcription and/or translation using a set of 221 publicly available metatranscriptome sets, comprising human gut (142), peat (48), plant (13), and symbionts (17); and 109 publicly available metaproteomes from 37 habitats - ***Table SI9***. Using bwa v.0.7.17^105^, AMP genes were mapped against the reads from the metatranscriptomes, and with NGLess^72^, we selected those genes with at least 1 read mapped across a minimum of two samples. Using Regex methods implemented in Python 3.8^76^, k-mers of all AMPSphere peptides (with length equal to at least half the length of the sequence) were checked for peptide sequences in metaproteomics data. In the case of perfect matches between a k-mer and a metaproteomic peptide for more than half the length of the sequence, it was considered that there is additional evidence that this c_AMP is likely to be expressed, as described by Ma et al.^5^. Briefly, the number of mapped peptides against the set of samples was counted and those peptides with at least 1 match covering more than 50% of the peptide were marked as detected. c_ AMPs with experimental evidence in metatranscriptomes and/or metaproteomes accounted for 1/5 of the AMPSphere.

We separated AMPs passing all quality-control tests into those with experimental evidence of translation/transcription (17,115 c_AMPs, ∼2% of AMPSphere) and those without it (63,098 c_AMPs, ∼7%). Quality filters for families consisted of keeping only those with ≥ 75% of its c_AMPs passing all quality control tests or having at least 1 c_AMP with experimental evidence of translation/transcription.

### Sample-based c_AMPs accumulation curves

For each habitat and group of habitats, we computed the sample-based accumulation curves by randomly sampling metagenomes 32 times in steps of 10 metagenomes. At each step, the number of unique c_AMPs found was computed, and the curves were drawn with the average obtained across the permutations.

### Multi-habitat and rare c_AMPs

The c_AMPs present simultaneously in ≥2 habitat groups were computed. To test the significance of this number, we opted for a similar approach to that described in Coelho et al. ^46^. The number of c_AMPs present in more than 1 habitat (high-level or general), here considered “multi-habitat”, was determined by counting the hits for each c_AMP in each sample. After that, the habitat labels for each sample were shuffled 100 times and the number of randomly obtained multi-habitat c_AMPs was counted. Shapiro-Wilks test was used to check the data distribution as normal (for general habitats - p = 0.49; and high-level habitats - p = 0.1) and this resulted in 676,489.7 ± 4281.8 multi-habitat c_AMPs by chance for high-level habitat groups, and in 685,477.17 ± 4,369.6 multi-habitat c_AMPs by chance for general habitats. High-level habitat groups presented 93,280 multi-habitat c_AMPs, while general habitats presented 173,955 multi-habitat c_AMPs. Both cases were 136.21 and 117.1 standard deviations below the value expected by chance, respectively. This was significant for the two cases with low estimated p-values (p < 10^−300^).

To determine the rarity of c_AMPs, the non-redundant genes in AMPSphere were mapped against the reads of metagenome samples using NGLess^72^. We considered only uniquely mapped reads. From the mapping, we computed the c_AMPs detected per sample and the number of detections per c_AMP, considering “rare”, the c_AMPs detected less than the average of the entire AMPSphere (682 detections or 1% of all samples). This approach was adopted to overcome the high computational costs involving a competitive mapping procedure. We expect our approach overestimates how prevalent the c_AMPs are, and because of that, it is a robust way to estimate the rarity observed in c_AMPs.

### Significance of the overlap of c_AMP contents

Similar to the significance testing of multi-habitat c_AMPs, the number of overlapping c_AMPs was computed for each pair of habitats (general and high-level). We shuffled the sample labels 1,000 times, counting the number of randomly overlapping c_AMPs for each pair of habitats, and used the Shapiro-Wilk test to verify normal distributions. Then, we estimated the probability of observing the overlap by Chebyshev’s inequality, which does not rely on normal distributions: 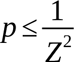, where Z stands for the Z-score computed from the average and standard deviations estimated by the shuffling procedure. The probabilities were adjusted using Holm-Sidak implemented in multipletests from the statsmodels package^106^, and those below 5% were considered significant.

### Differences in the c_AMP density in microbial species from different habitats

The c_AMP density was defined as *ρ_AMPs_=n_C_AMPs__/L*, where *n_C_AMPs__* is the number of c_AMP redundant genes and L is the assembled base pairs. It was computed per sample to verify the differences between animal host-and non-host-associated samples, including only habitats with ≥100 samples. The calculated densities were filtered by using Tukey’s fences calculated with *k* =1.5 to eliminate outliers.

This density was also calculated at phylum, species, and genus levels, summing all assembled base pairs for contigs assigned to each one of those taxonomy levels in the samples used in AMPSphere. In that case, we assume, as an approximation, that in a large assembled segment, the start positions of AMP genes are independent and uniformly random. Thus, we calculated the standard sample proportion error, with the formula: 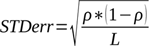. The error was used to calculate the margin of error at a 95% confidence interval (*Z*=1.96,). Genera, phyla and species *ρ_AMP_* within a margin of error superior to 10% of the calculated value were eliminated along with outliers according to Tukey’s fences (*k* =1.5). To verify the effects of different habitats in the *ρ_AMP_* of species, we took the density calculated per species per sample from species present in ≥2 habitats in ≥10 samples per habitat and tested their medians using the Mann-Whitney U test implemented in the scipy package^80^.

We verified the species abundances in each sample using mOTUs 2^107^. None of the genera found as those with the highest *ρ_AMP_* (*Algorimicrobium*, TMED78, SFJ001, STGJ01, and CAG-462) were also verified throughout the mOTUs, which show them as not abundant microbes.

To evaluate the effects of potential fragments in our density analysis, all tests and comparisons were also performed restricting the set of c_AMPs used to only those with at least one stop codon upstream the gene. The results showed that results related to the c_AMP density observed for all AMPSphere did not modify when controlling for quality (***Fig. SI5***).

Differences between habitats for each species, genus, or sample groups were tested using Mann-Whitney U and Kruskal-Wallis tests implemented in the scipy package^80^. P-values were adjusted using the Holm-Sidak method implemented in multipletests from the statsmodels package^106^.

### Determination of accessory AMPs

Core, shell, and accessory c AMP clusters were determined using the subset of c_AMPs obtained from ProGenomes v2^41^ because of their high-confidence assigned taxonomies and the genomically-defined species (specI). To increase confidence in our measures, only species containing ≥10 genomes were considered. AMPs and families (^≥^8 c_AMPs) present in fewer than 50% of the genomes from a microbial species were classified as accessory. Those c_AMPs and families present in 50% - 95% of the genomes in the cluster were classified as shell^108^, and those present in >95% of the genomes were classified as core genes^51^.

We used FastANI v.1.33^88^ to cluster genomes in species in the ProGenomes v2^41^, keeping one randomly selected representative for each clonal complex (ANI ≥ 99.99%) and inferring strains (99.5% ≤ ANI < 99.99%) as in Rodriguez et al.^109^. Only species with ≥10 genomes after elimination of clonal redundancy were kept. To verify the propensity of AMPs being shared between genomes belonging to the same strains, we computed all possible pairs of genomes per species and computed the pairs sharing AMPs, testing the results with the Fisher’s Exact test implemented in the scipy package^80^. We also extracted the predicted full-length proteins from the ENA database for each genome and hierarchically clustered them after alphabet reduction in a similar fashion to that described in the topic “***AMP families***”, keeping those clusters with ≥8 sequences for each species. The prevalence of full-length protein families within a species was computed as above mentioned and the number of core families was compared to the number of c_AMP core families using the probability calculated as number of species with proportion of core full-length protein families less or equal to that observed for c_AMPs divided by the total of assessed species.

To determine the genotype of *Mycoplasma pneumoniae* samples in ProGenomes2^41^, the gene coding for P1 adhesin^53^ was mapped against the genomes using the reference gene NZ_LR214945.1:c568695-567307 with bwa^105^, and later extracted with SAMtools^110^ and BEDtools^111^. The extracted genes were aligned using Clustal Omega^112^, and a phylogenetic tree was built using nucleotide sequences and FastTree 2^87^ with the restricted time-reversible substitution model and a bootstrapping procedure with 1,000 pseudo-replicates to determine node support. The tree was used to segregate and classify genomes taking the strain type of reference genomes from Diaz et al.^54^ and was consistent with the previously established groups.

### Annotation of AMPs using different datasets

Databases used in the annotation were the small protein sets in SmProt 2^45^, the bioactive peptides database starPepDB 45k^91^, the small proteins from the global data-driven census of *Salmonella*^113^, the global microbial gene catalog GMGCv1^46^, and a specific AMP database – DRAMP 3.0^44^. To only have sequences that were unlikely to be artifacts of assembly for the analysis, only c_AMPs passing the terminal placement test were searched against the GMGCv1^46^. The AMPs were annotated using MMseqs2^75^ with the ‘easy-search’ method, retaining hits with an E-value maximum of 10^−5^. To normalize coordinates of hits to the full-length protein, we corrected for the elimination of the initial methionine performed by Macrel^40^, so that hits starting at the second amino acid were considered as if they matched the first one (as the peptide has had its initial methionine removed).

We used the hypergeometric test implemented in the scipy package^80^ to model the association between c_AMPs and the background distribution of ortholog groups from GMGCv1^46^. To that, the number of genes in the redundant GMGCv1^46^ for each ortholog group was computed along with the counts for ortholog groups in the top hits to AMPSphere. The enrichment was given as the proportion of hits presenting a given ortholog group divided by the proportion of that ortholog group among the redundant sequences in GMGCv1^46^ and was considered significant when p < 0.05 after a correction with Holm-Sidak method implemented in multipletests from the statsmodels package^106^. With a robust approach, filtering the OGs by the number of c_AMP hits and GMGCv1^46^ hits associated with them, using a minimum of 10, 20, or even 100 proteins, the results were kept similar to those obtained with all data.

To check for genomic entities generated after gene truncation, we screened for c_AMP homologs using the default settings for Blastn^103^ against the NCBI database^94^, keeping only significant hits with a maximum E-value of 10^−5^. We selected the AMP10.271_016, predicted to be produced by *Prevotella jejuni*, which shares the start codon with the gene coding for a NAD(P)-dependent dehydrogenase (WP_089365220.1). Using Biopython^83^, we codon-aligned the fragments from metagenomic contigs assembled from samples SAMN09837386, SAMN09837387, and SAMN09837388, and genomic fragments of different strains of *Prevotella jejuni* CD3:33 (CP023864.1:504836-504949), F0106 (CP072366.1:781389-781502), F0697 (CP072364.1:1466323-1466436), and from *Prevotella melaninogenica* strains FDAARGOS_760 (CP054010.1:157726- 157839), FDAARGOS_306 (CP022041.2:943522-943635), FDAARGOS_1566 (CP085943.1:1102942-1103055), and ATCC 25845 (CP002123.1:409656-409769).

### Positive selection tests

The genes of c_AMPs belonging to 100 high-quality families randomly sampled were codon-aligned, excluding identical sequences and the stop codons. Selection tests were run using HyPhy version 2.5.1 (www.hyphy.org)^114^ on the codon alignments and trees obtained as previously mentioned. Alignment-wide episodic diversification tests were computed on the gene family clusters with the BUSTED method^115^. Genes with Holm-Bonferroni multiple-test corrected p-values lower than 0.05 were considered as positively selected. Only a small fraction (∼15%) of the tested families presented evidence of episodic diversification.

### Genomic context conservation analysis

We mapped the 863,498 AMP sequences against a collection of 169,632 reference genomes, MAGs and SAGs curated elsewhere^49^ with DIAMOND^116^ in blastp mode. Hits with identity > 50% (amino acid) and query and target coverage > 90% were considered significant. A total of 107,308 AMPs have homologs in at least one genome. We built gene families from the hits of each AMP detected in the prokaryotic genomes, and calculated a conservation score of the functional annotation of the neighboring genes in a window of 3 genes up and downstream. The vertical conservation score of each OG, KEGG pathway, KEGG orthology, KEGG module – from the Kyoto Encyclopedia of Genes and Genomes (KEGG)^117^, PFAM 33.1^92,97^, and CARD^118^ at each position was calculated as the number of genes with a given functional annotation divided by the number of genes in the family. AMPs with more than 2 hits and a vertical conservation score > 0.9 with any functional term were considered to have conserved genomic contexts.

For testing whether the number of AMPs with conserved genomic neighbors is higher than in other gene families within the 169,632 genomes curated by del Río et al.^49^, we used MMSeqs2^75^ for building *de novo* gene families (establishing the minimum amino acid identity of 30%, coverage of the shorter sequence of at least 50%, and maximum E-value of 10^−3^). We then took 10,000 random sets of 55,191 gene families with more than 2 members (number of AMPs with more than two homologs in the genomes) composed of: (i) small (< 50 residues) proteins, and (ii) no length-restricted proteins. Then, we computed the number of gene families showing conserved genomic contexts with known functions within each set and confirmed their normal distribution using the Shapiro-Wilks test^80^. Later, the conservation values obtained for AMPSphere were compared to those sets using P-values computed from their respective Z-scores.

To verify the ortholog groups from c_AMPs and the gene neighborhood, the peptides were also annotated using EggNOG-mapper v2^84^ using default settings and selecting the best hit for each c_AMP, by filtering the lowest E-value and highest best score. Their KEGG ortholog groups (KOs) were used as functional labels to cluster and verify the gene neighborhood in terms of functional similarity. It was possible to annotate 56.1% (60,173 out of 107,308) of c_AMPs with hits to the genome set tested using the EggNOG5 database^48^, with 9.1% of them missing COG categories, and about 18.1% of them belonging to translation-related functions (J), 14.4% belonging to unknown function proteins (S), and 9% of them belonging to replication, recombination, and repair (L).

### c_AMPs and bacterial species transmissibility

We used the species taxonomy and transmissibility indexes calculated by Valles-Colomer et al.^57^ to demonstrate the effect of AMPs on the transmission of bacterial species from mother to children. Only those species overlapping AMPSphere and the datasets from Valles-Colomer et al.^57^ were kept for this analysis, and their AMP densities were calculated separately for samples from the human gut and human oral cavity as mentioned in the section ***Differences in the c_AMP density in microbial species from different habitats***. The AMP density and the coefficient of transmissibility were correlated using Spearman’s method implemented in the scipy package^80^ to keep robust comparisons and were tested for different situations, e.g. following children’s microbiome after 1, 3, and up to 18 years, as well as, cohabitation and intra-datasets. Missing data was completely omitted for calculus purposes.

### AMPSphere web resource

AMPSphere is found at the address https://ampsphere.big-data-biology.org/. The implementation is based on Python^76^ and Vue Javascript. The database was built with sqlite, and SQLalchemy was used to map the database to Python objects. Internal and external APIs were built using FastAPI and Gunicorn to serve them. In the front end, Vue 3 was used as the backbone and Quasar built the layout. Plotly was used to generate interactive visualization plots, and Axios to render content seamlessly. LogoJS (https://logojs.wenglab.org/app/) is used to generate sequence logos for AMP families; while the helical wheel app (https://github.com/clemlab/helicalwheel) generates AMP helical wheels.

### Selection of peptides to synthesis and activity testing

Only high-quality (see the topic “***Quality control of c_AMPs***”) c_AMPs were considered for synthesis. They were filtered according to 6 criteria for solubility and 3 criteria for synthesis, as used before in PepFun^119^. A peptide approved for at least 6 of these criteria was then filtered by predicting AMP activity with 6 methods in addition to Macrel^40^: AMPScanner v2^120^, ampir^38^, amPEPpy^121^, APIN^122^, and AMPLify^123^. Peptides predicted to be AMPs by all methods were filtered by length, discarding sequences longer than 40 amino acid residues, for which conventional solid-phase peptide synthesis using Fmoc strategy has lower yields and many recoupling reactions^124–127^. AMPs were sorted by their abundance (the number of redundant genes), keeping the most abundant peptide per AMP family. After this process, we obtained 364 candidate AMPs, belonging to 166 families and 198 clusters with <8 c_AMPs. Of those, 30 candidates were homologous to sequences from the databases used in annotation (*e.g.,* SmProt 2^45^). To compose the list of 50 peptides proceeding to synthesis and testing: (i) we selected 34 of the most abundant peptides, eliminating peptides that overlap with other candidates and replacing them with the next most abundant ones; (ii) randomly selecting c_AMPs with homologs to a final proportion of 30% of our set, more specifically 14 peptides that matched GMGCv1^46^, and 1 that matched SmProt 2^45^; and (iii) 1 peptide also found in the MAGs binned from stool samples used to investigate fecal transplantations^128^.

### Minimal inhibitory concentration determination

The 50 AMPs were tested for antimicrobial activity using the broth microdilution method^129^. MIC values were considered as the concentration of the peptides that killed 100% of cells after 24 h of incubation at 37°C. First, peptides diluted in water were added to untreated flat-bottom polystyrene microtiter 96-well plates in two-fold dilutions ranging from 64 to 1 μmol·L^−1^, and then peptides were exposed to an inoculum of 2·10^6^ cells in LB or BHI broth, for pathogens and gut commensals, respectively. After the incubation time, the absorbance of each well representing each of the conditions was analyzed using a spectrophotometer at 600 nm. The assays were conducted in three biological replicates to ensure statistical reliability.

### Circular dichroism assays

Circular dichroism experiments were conducted using a J1500 circular dichroism spectropolarimeter (Jasco) at the Biological Chemistry Resource Center (BCRC) of the University of Pennsylvania. The experiments were carried out at a temperature of 25°C. Circular dichroism spectra were obtained by averaging three accumulations using a quartz cuvette with an optical path length of 1.0 mm. The spectra were recorded in the wavelength range from 260 to 190 nm at a scanning rate of 50 nm·min^−1^ with a bandwidth of 0.5 nm. The peptides were tested at a concentration of 50 μmol·L^−1^. Measurements were performed in water, a mixture of water and trifluoroethanol (TFE) in a ratio of 3:2, and a mixture of water and methanol in a ratio of 1:1. Baseline measurements were recorded prior to each measurement. To minimize background effects, a Fourier transform filter was applied. The helical fraction values were calculated using the single spectra analysis tool available on the BeStSel server^130^.

### Outer membrane permeabilization assays

Membrane permeability was analyzed using the 1-(N-phenylamino)naphthalene (NPN) uptake assay. NPN demonstrates weak fluorescence in an extracellular environment but displays strong fluorescence when in contact with lipids from the bacterial outer membrane. Thus, NPN will show increased fluorescence when the integrity of the outer membrane is compromised. *A. baumannii* ATCC 19606 and *P. aeruginosa* PA01 were cultured until cell numbers reached an OD_600_ of 0.4, followed by centrifugation (10,000 rpm at 4°C for 3 min), washing, and resuspension in buffer (5 mmol·L^−1^ HEPES, 5 mmol·L^−1^ glucose, pH 7.4). Subsequently, 4 µL of NPN solution (working concentration of 0.5 mmol·L^−1^) was added to 100 µL of bacterial solution in a white flat bottom 96-well plate. The fluorescence was monitored at λ_ex_ = 350 nm and λ_em_ = 420 nm. The peptide solutions in water (100 µL solution at their MIC values) were introduced into each well, and fluorescence was monitored as a function of time until no further increase in fluorescence was observed (30 min). The relative fluorescence was calculated using a non-linear fit. The positive control (antibiotic polymyxin B) was used as baseline. The following equation was applied to reflect % of difference between the baseline (polymyxin B) and the sample:

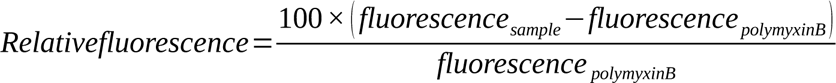

### Cytoplasmic membrane depolarization assays

The ability of the peptides to depolarize the cytoplasmic membrane was assessed by measuring the fluorescence of the membrane potential-sensitive dye 3,3’-dipropylthiadicarbocyanine iodide [DiSC_3_-(5)]. This potentiometric fluorophore fluoresces upon release from the interior of the cytoplasmic membrane in response to an imbalance of its transmembrane potential. *A. baumannii* ATCC 19606 and *P. aeruginosa* PA01 cells were grown with agitation at 37°C until they reached mid-log phase (OD_600_ = 0.5). The cells were then centrifuged and washed twice with washing buffer (20 mmol·L^−1^ glucose, 5 mmol·L^−1^ HEPES, pH 7.2) and re-suspended to an OD_600_ of 0.05 in 20 mmol·L^−1^ glucose, 5 mmol·L^−1^ HEPES, 0.1 mol·L^−1^ KCl, pH 7.2. An aliquot of 100 µL of bacterial cells was added to a black flat bottom 96-well plate and incubated with 20 nmol·L^−1^ of DiSC_3_-(5) for 15 min until the fluorescence stabilized, indicating the incorporation of the dye into the cytoplasmic membrane. The membrane depolarization was monitored by observing the change in the fluorescence emission intensity of the dye (λ_ex_ = 622 nm, λ_em_ = 670 nm), after the addition of the peptides (100 µL solution at their MIC values). The relative fluorescence was calculated using a non-linear fit. The positive control (antibiotic polymyxin B) was used as baseline. We estimated the % of difference between the baseline (polymyxin B) and the sample using the same mathematical approach as in the “***Outer membrane permeabilization assays***”.

## QUANTIFICATION AND STATISTICAL ANALYSIS

Graphs for the experimental results were created and statistical tests conducted in GraphPad Prism v.9.5.1 (GraphPad Software, San Diego, California USA).

## ADDITIONAL RESOURCES

AMPSphere is freely available for download in Zenodo^131^ and as a web server (https://ampsphere.big-data-biology.org/). The code related to the AMPSphere generation and analysis is available in: https://github.com/BigDataBiology/AMPsphere.

## Supplemental Information

**Figure SI1.**
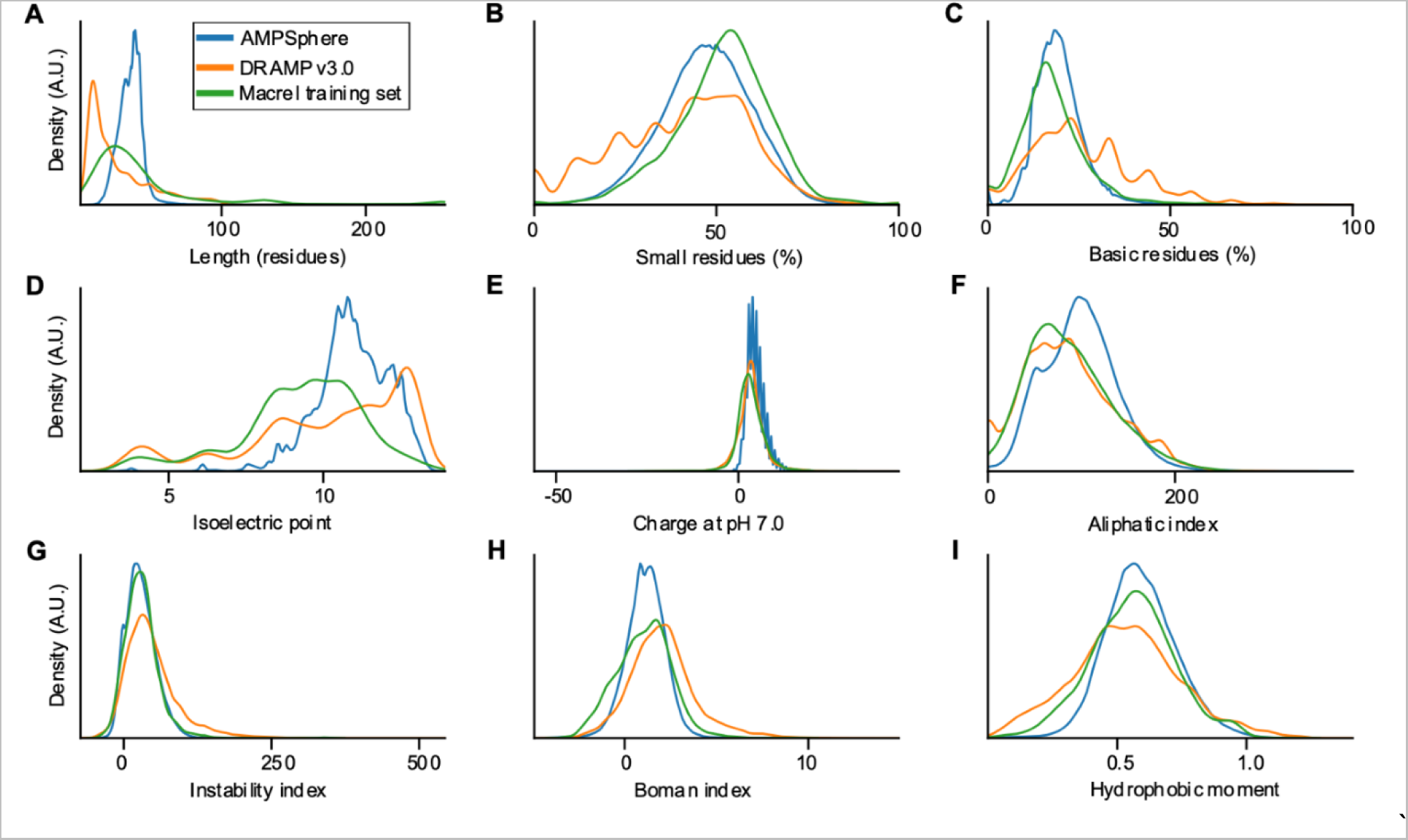
General physical-chemical features of c_AMPs in AMPSphere and validated databases of antimicrobial peptides. All graphs are drawn using the continuous probability density curve and given as functions of the arbitrary density units. The c_AMPs from AMPSphere retain the same predicted biochemical properties of those from validated AMPs, although with an average length longer than the peptides in the training set of Macrel [S1] and DRAMP 3.0 [S2]. (***A***) The distribution of peptide length in residues across the AMPSphere. (***B***) Distribution of proportions of residues with small side chains [A, B, C, D, G, N, P, S, T, V] per AMP. (***C***) Distribution of proportions of basic residues [H, R, K] per AMP. (***D***) Distribution of isoelectric points in AMPSphere peptides. (***E***) Distribution of peptide charges at pH 7.0. (***F***) Distribution of aliphatic index in peptides from AMPSphere. (***G***) Distribution of instability index in AMPSphere. (***H***) Distribution of Boman index in AMPSphere. (***I***) Distribution of hydrophobic moments in AMPSphere.

**Figure SI2.**
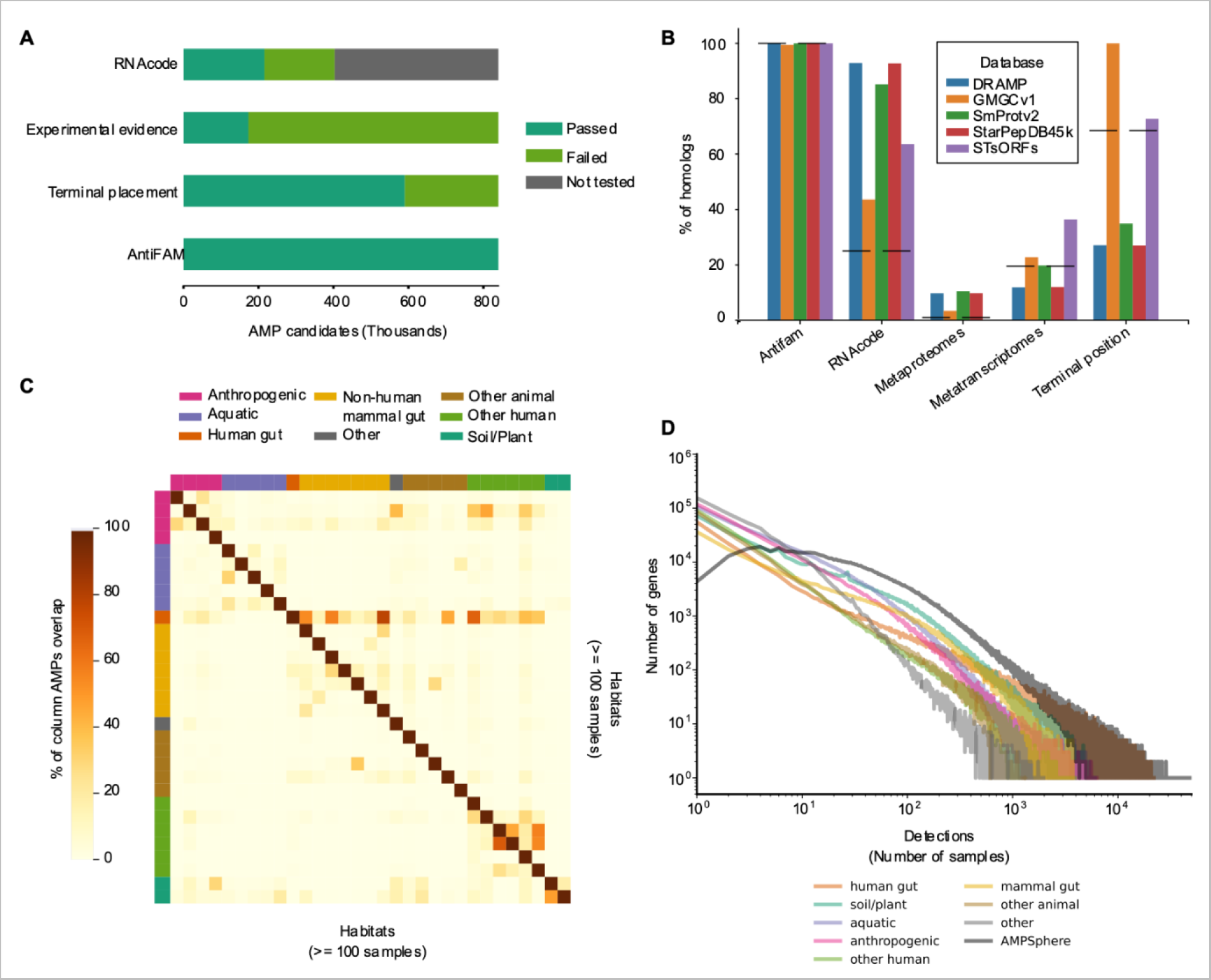
c_AMPs quality and habitat distribution. (***A***) Quality assessment of AMPSphere reveals most of the peptides passing at least 1 of the tests. The RNAcode test depends on gene diversity, which is very low for AMPSphere, and therefore, determines a low rate of positives among our candidates. (***B***) c_AMPs homologous to databases of validated bioactive peptides also showed a higher average quality of these datasets. (***C***) The limited overlap of c_AMPs among habitats argues in favor of using habitat groups to gain resolution. Note that the group of habitats with the highest paired overlaps belong to human body sites and samples from human guts and non-human mammalian guts. Only habitats with at least 100 samples were shown. (***D***) It is also possible to observe the great proportion of rare genes in AMPSphere from different habitat groups, in which few genes are largely detected.

**Figure SI3.**
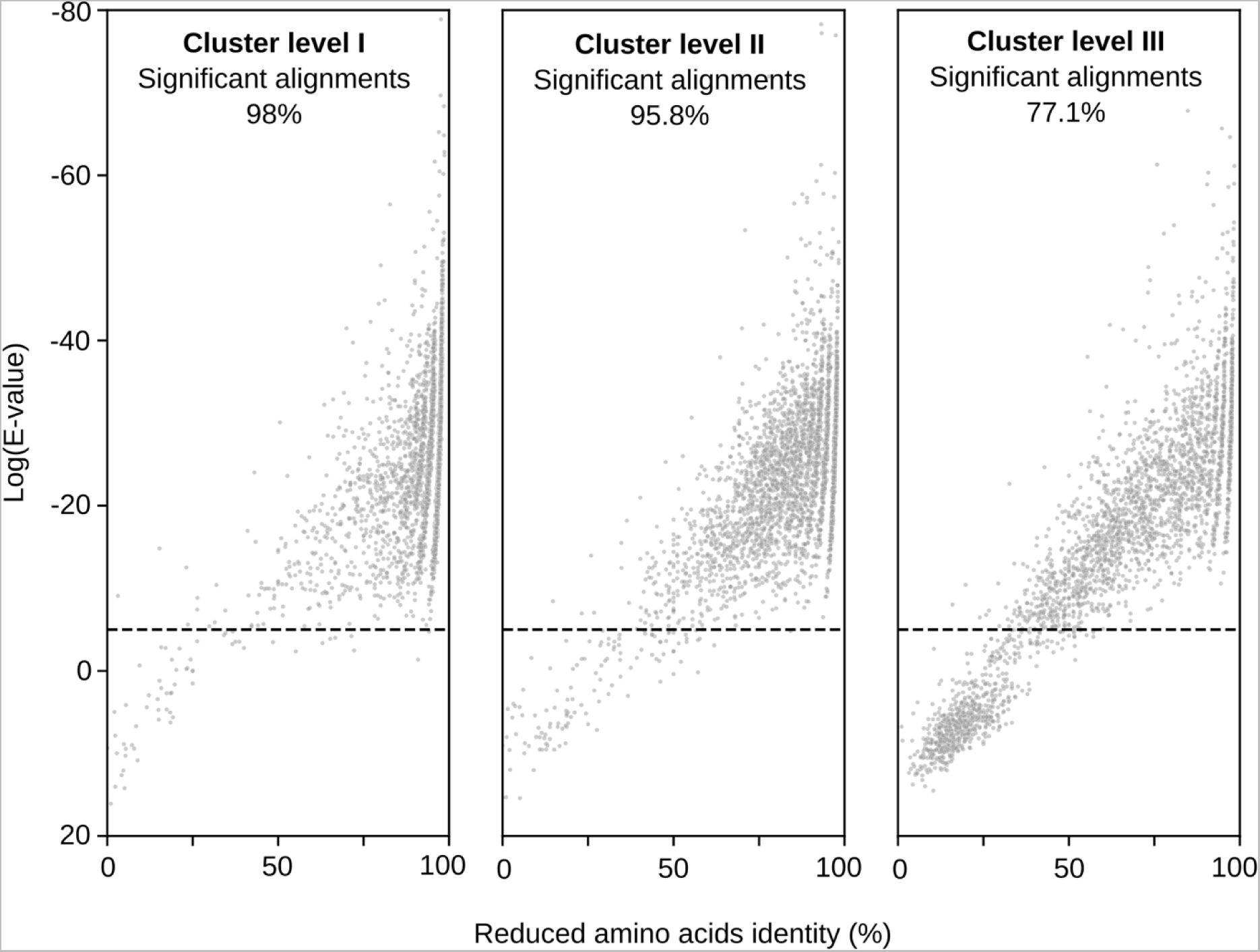
Clustering validation of families. To validate the clustering procedure using a reduced amino acid alphabet, samples of 1,000 peptides were randomly drawn from AMPSphere (excluding representative sequences) and aligned against their cluster representatives. Three different levels (I, II, and III) of clustering were tested. The E-values were computed per alignment and plotted against the corresponding alignment identity. The averaged proportion of significant alignments is shown above each graph.

**Figure SI4.**
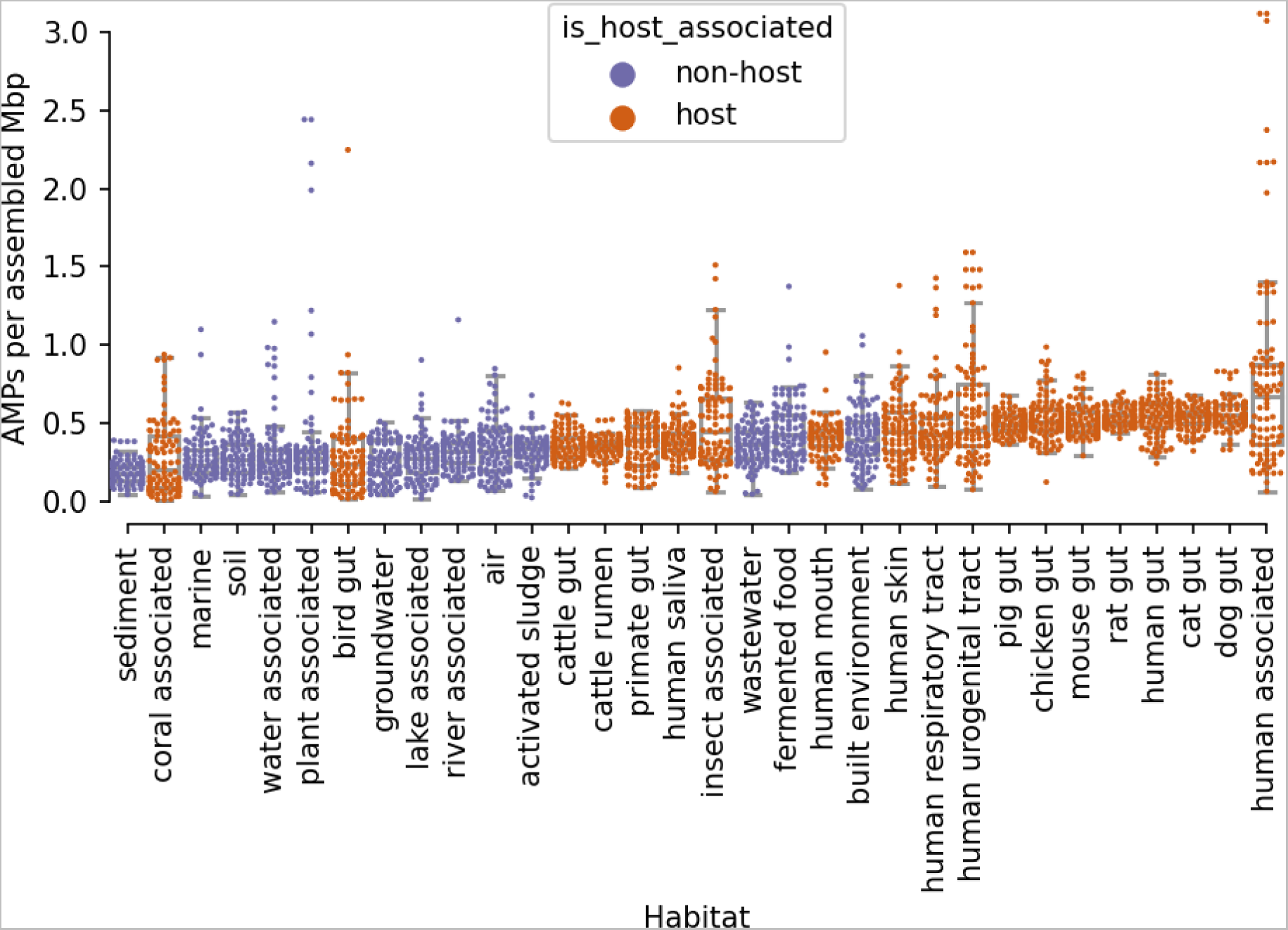
Host-associated habitats are denser in c_AMPs. The c_AMP density (given as c_AMP genes per assembled Mbp) was computed per sample and plotted per habitat. Only habitats with at least 100 samples were used. To favor a good visualization, we sampled 2,000 dots to plot their distribution trends. habitats were colored by their ontology, orange for the animal host-associated samples and purple for the non-animal-host-associated samples. Host-associated habitats cluster at the right portion of the graph with some anthropogenic habitats, which are closer to animal host-associated samples than habitat samples. Few exceptions, such as coral-associated and bird gut habitats cluster with non-animal-host associated habitats.

**Figure SI5.**
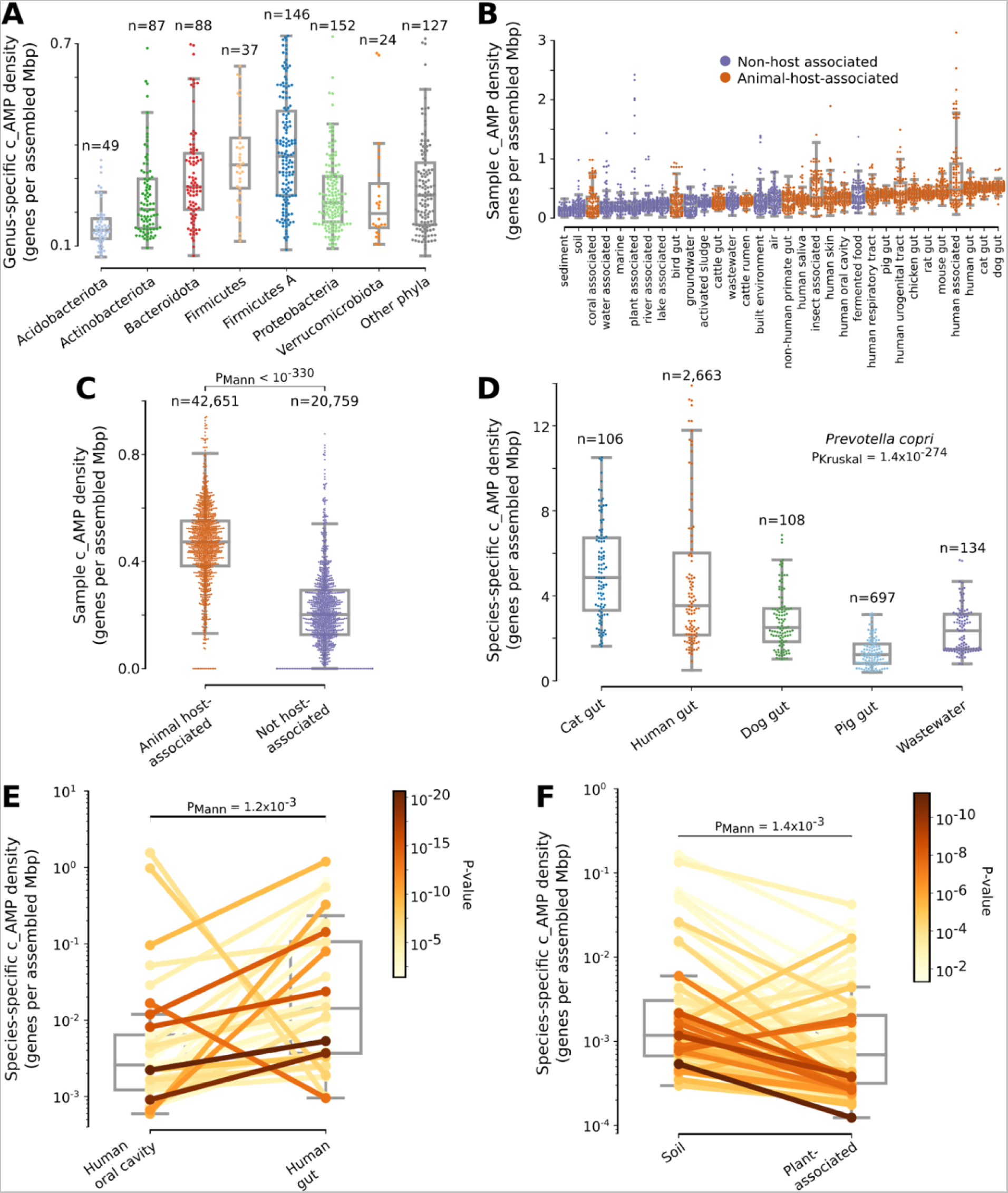
Quality-controlled c_AMPs do not change the differences observed for c_AMP density across several bases of comparison. To evaluate the effects of possible gene fragments, we filtered off genes not having a stop codon upstream of them and recalculated the c_AMP density (given as c_AMP genes per assembled Mbp). The same results could be observed in the correspondent analysis using all c_AMPs. ***(A)*** The main phyla contributing with more c_AMPs in AMPSphere and the distribution of these densities in each of those groups by genera – correspondent to Fig. 4C. ***(B)*** Animal host-associated habitats are denser in c_AMPs - correspondent to ***Fig. SI4. (C)*** Animal host-associated habitats have a higher sample c_AMP density when compared to those non-host-associated - correspondent to Fig. 5A. ***(D)*** Hosts are a factor for variation of c_AMP density in *Prevotella copri*, presenting a higher ⍴AMP in cat and human guts compared to the same species in guts of pigs and dogs - correspondent to Fig. 5B. 106 randomly selected points are shown for each host. ***(E)*** Species-specific ⍴*_AMP_* of microbes from the human gut are higher, when compared against the same species found in the human oral cavity - correspondent to Fig. 5C. ***(F)*** For non-animal hosts, the species-specific ⍴*_AMP_* of microbes from the soil is higher when compared against the same species found in plant-associated samples - correspondent to Fig. 5D. For panels ***E*** and ***F*** the significance was color-encoded using a Log_10_(P_Mann_) scale.

**Figure SI6.**
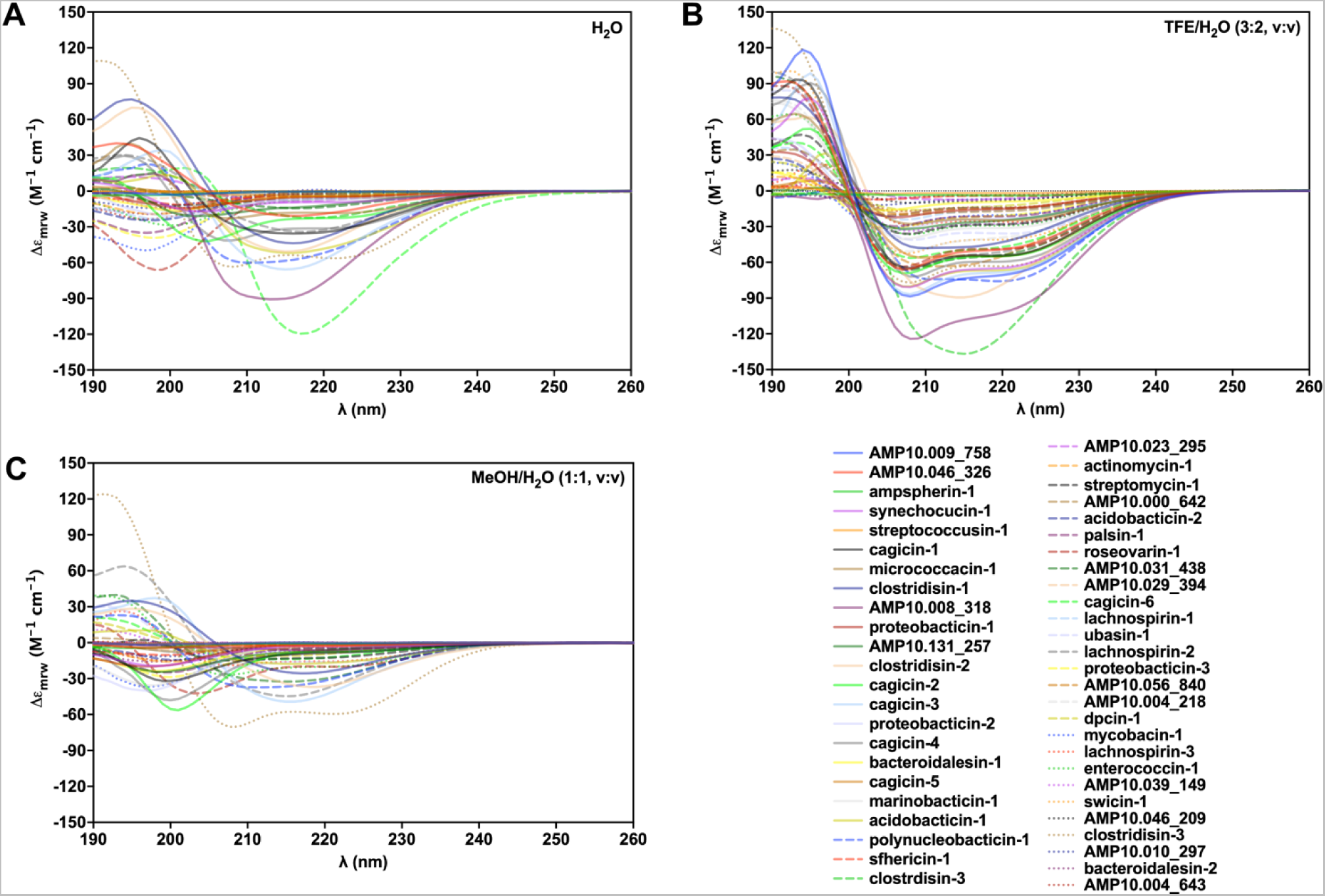
Circular dichroism spectra of the c_AMPs. c_AMPs secondary structural tendency was analyzed using three different solvents: ***(A)*** water, ***(B)*** trifluoroethanol (TFE) and water mixture (3:2, V:V), and ***(C)*** methanol (MeOH) and water mixture (1:1, V:V). The experiments were carried out at 25 °C, and the circular dichroism spectra shown are an average of three accumulations obtained using a quartz cuvette with an optical path length of 1.0 mm, ranging from 260 to 190 nm at a rate of 50 nm·min^−1^ and a bandwidth of 0.5 nm. All peptides were tested at a concentration of 50 μmol·L^−1^, with respective baselines recorded prior to measurement. A Fourier transform filter was applied to minimize background effects.

**Figure SI7.**
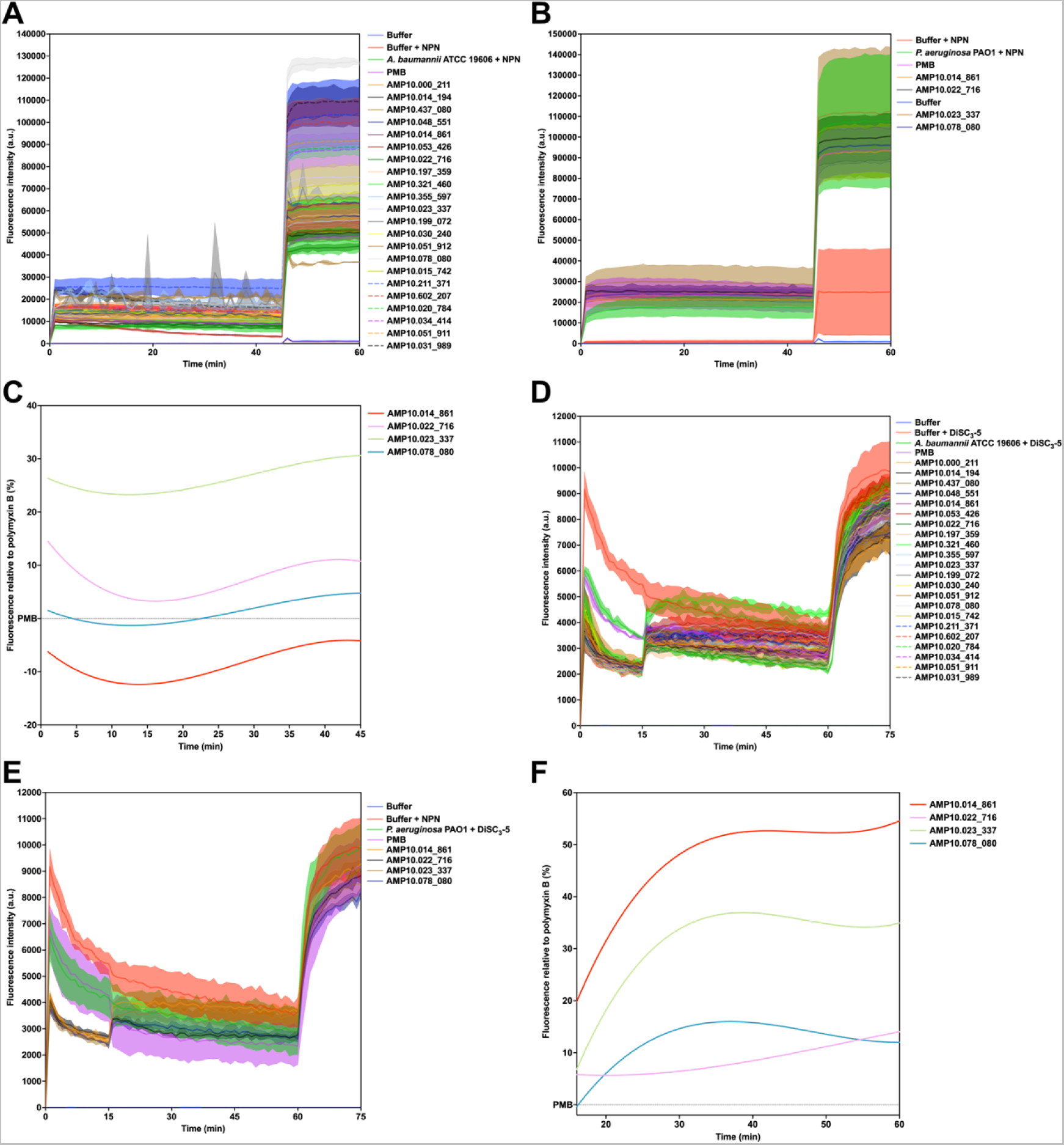
Mechanism of action of AMPSphere peptides. Permeabilization assays with the fluorescent probe 1-(N-phenylamino)naphthalene (NPN) showing the effect of AMPSphere peptides on ***(A)*** *A. baumannii* ATCC 19606 and ***(B)*** *P. aeruginosa* PA01 cells, and ***(C)*** Fluorescence values relative to polymyxin B (PMB, positive control) of the fluorescent probe 1-(N-phenylamino)naphthalene (NPN) that indicate outer membrane permeabilization of *P. aeruginosa* PA01 cells. Depolarization assays with the hydrophobic probe 3,3′-dipropylthiadicarbocyanine iodide [DiSC3-(5)], effect of AMPSphere peptides in ***(D)*** *A. baumannii* ATCC 19606 and ***(E)*** *P. aeruginosa* PA01 cells. ***(F)*** Fluorescence values relative to PMB (positive control) of 3,3′-dipropylthiadicarbocyanine iodide [DiSC3-(5)], a hydrophobic fluorescent probe, used to indicate cytoplasmic membrane depolarization of *P. aeruginosa* PA01 cells. Data in ***(A)***, ***(B)***, ***(D)***, and ***(E)*** are the mean plus and minus the standard deviation.

## Excel Tables

***Table SI1. Metadata and description of (meta)genomes used in AMPSphere.*** The sample is identified by its access code in ENA, the habitat shows the type of habitat this sample was retrieved from. Other data about the sequencing, such as the number of raw inserts and the number of assembled base pairs (bp) are also available along with the information on N50. The number of predicted complete large ORFs (>100 amino acids) and smORFs (10-100 amino acids) is shown (ORFs+smORFs) along with the number of smORFs alone and the predicted non-redundant c_AMPs.

***Table SI2. c_AMP distribution in the habitat groups.*** The habitats grouped under each class are shown along with the number of genes encoding the non-redundant c_AMPs, the number of c_AMP clusters in total, and the number of clusters containing ≥8 c_AMPs (c_AMP families).

***Table SI3. Ortholog groups (OGs) enrichment in the hits to the GMGCv1*** [S3]. Top hits were assessed and the proportion of OGs from eggNOG 5 [S4] was compared using the number of c_AMPs affiliating to homologs of a given OG and the total number of OGs found in the homologs of c_AMPs (156,711) in the comparison to the GMGCv1 [S3]. As a background measure, we used the counts of a given OG in the redundant set of genes belonging to GMGCv1 [S3] and the total number of OGs found in the redundant GMGC catalog [S3] (9,180,087,363). Enrichment in the c_AMPs set was given as the fold-change calculated for each given OG in relation to that expected in the GMGCv1 [S3]. P-values were adjusted using Holm-Sidak and only significant hits (P < 0.05) were shown.

***Table SI4. c_AMP genome context in comparison to families with proteins of different sizes.*** The proportion of families of proteins of different sizes (all lengths and only ≤ 50 amino acids) is presented in comparison to the proportion of mapped AMPs (55,191) with genome contexts involving a given Kyoto Encyclopedia of Genes and Genomes – KEGG ortholog pathway [S5] shown with their accession code and description.

***Table SI5. Permutations with random families of different sizes show that the genome context of c_AMPs is different from other protein families.*** We performed 10,000 random samplings of 55,191 protein families of all lengths and only smaller than 50 amino acids. These families were assessed regarding their genome neighborhood conservation. It was found that the values found in AMPSphere are mostly different from those observed for any other protein families across genome contexts involving known functions, antibiotics synthesis/resistance, and antibiotic resistance genes from CARD [S6].

***Table SI6. AMPs with conserved genome contexts sharing KEGG ortholog groups (KO) with gene neighbors.*** AMPs were annotated with eggNOG mapper [S7] and those harboring KOs were included in this analysis. It is shown for each KO a brief description of its activity, the total number of AMPs annotated, the number of those with conserved neighbors (conservation score > 0.9), and the number of AMPs assigned to a given KO inserted in a conserved neighborhood containing other genes annotated to that KO (the intersection before the 2 previous columns).

***Table SI7. c_AMP density across different genera.*** The number of redundant c_AMP genes as well as their respective assembled base pairs is presented with the calculated AMP density. The error on the AMP density measure is also shown evidence that most of the genera present an error above 10% (our cutoff).

***Table SI8. Differential species-specific c_AMP density across habitats.*** The tested species had their densities (as c_AMP genes per assembled gigabase pairs) compared across samples from different habitats (Habitats A and B), and the number of samples for each habitat was also registered after eliminating the outliers using Tukey’s fences with k = 1.5 (# Samples A and B). The average c_AMP density for each species in each habitat was registered along with its standard deviation (Avg. c_AMP density and Std. c_AMP density). The Mann-Whitney U test was applied to each pair of habitats and later corrected using the Holm-Sidak method, shown as the ‘Adjusted P-value’. Comparisons considered species present in at least 10 samples for each habitat with ≥ 100 samples presenting c_AMPs. Only significant comparisons were shown.

***Table SI9. Metatranscriptomes and metaproteomes used in the verification for experimental signals of transcription and/or translation of c_AMP genes from AMPSphere.*** Metatranscriptomes from EMBL-ENA were used in the comparisons with genes in AMPSphere to verify signals of transcription in datasets *ad hoc*. The datasets from the Proteomics Identification Database (PRIDE) - EMBL-EBI were also used in the comparison to c_AMPs and verified the same peptides in datasets *ad hoc*.

## Notes

### Summary of Updates

Author names had first and last names reversed.

## REFERENCES

1. Antimicrobial Resistance Collaborators (2022). Global burden of bacterial antimicrobial resistance in 2019: a systematic analysis. Lancet 399, 629–655. 10.1016/S0140-6736(21)02724-0.

2. Stokes, J.M., Yang, K., Swanson, K., Jin, W., Cubillos-Ruiz, A., Donghia, N.M., MacNair, C.R., French, S., Carfrae, L.A., Bloom-Ackermann, Z., et al. (2020). A Deep Learning Approach to Antibiotic Discovery. Cell 180, 688–702.e13. 10.1016/j.cell.2020.01.021.

3. Torres, M.D.T., Melo, M.C.R., Flowers, L., Crescenzi, O., Notomista, E., and de la Fuente-Nunez, C. (2022). Mining for encrypted peptide antibiotics in the human proteome. Nat Biomed Eng 6, 67–75. 10.1038/s41551-021-00801-1.

4. Porto, W.F., Irazazabal, L., Alves, E.S.F., Ribeiro, S.M., Matos, C.O., Pires, Á.S., Fensterseifer, I.C.M., Miranda, V.J., Haney, E.F., Humblot, V., et al. (2018). In silico optimization of a guava antimicrobial peptide enables combinatorial exploration for peptide design. Nat Commun 9, 1490. 10.1038/s41467-018-03746-3.

5. Ma, Y., Guo, Z., Xia, B., Zhang, Y., Liu, X., Yu, Y., Tang, N., Tong, X., Wang, M., Ye, X., et al. (2022). Identification of antimicrobial peptides from the human gut microbiome using deep learning. Nat Biotechnol, 1–11. 10.1038/s41587-022-01226-0.

6. Besse, A. (2017). Halocin C8: an antimicrobial peptide distributed among four halophilic archaeal genera: Natrinema, Haloterrigena, Haloferax, and Halobacterium. Extremophiles 21. 10.1007/s00792-017-0931-5.

7. Cotter, P.D., Ross, R.P., and Hill, C. (2013). Bacteriocins — a viable alternative to antibiotics? Nat Rev Microbiol 11, 95–105. 10.1038/nrmicro2937.

8. Wang, S., Zheng, Z., Zou, H., Li, N., and Wu, M. (2019). Characterization of the secondary metabolite biosynthetic gene clusters in archaea. Comput Biol Chem 78, 165–169. 10.1016/j.compbiolchem.2018.11.019.

9. Zasloff, M. (2019). Antimicrobial Peptides of Multicellular Organisms: My Perspective. In Antimicrobial Peptides: Basics for Clinical Application, K. Matsuzaki, ed. (Springer Singapore), pp. 3–6. 10.1007/978-981-13-3588-4_1.

10. Huang, K.-Y., Chang, T.-H., Jhong, J.-H., Chi, Y.-H., Li, W.-C., Chan, C.-L., Robert Lai, K., and Lee, T.-Y. (2017). Identification of natural antimicrobial peptides from bacteria through metagenomic and metatranscriptomic analysis of high-throughput transcriptome data of Taiwanese oolong teas. BMC Syst Biol 11. 10.1186/s12918-017-0503-4.

11. Heilbronner, S., Krismer, B., Brötz-Oesterhelt, H., and Peschel, A. (2021). The microbiome-shaping roles of bacteriocins. Nat Rev Microbiol, 1–14. 10.1038/s41579-021-00569-w.

12. Pizzo, E., Cafaro, V., Di Donato, A., and Notomista, E. (2018). Cryptic Antimicrobial Peptides: Identification Methods and Current Knowledge of their Immunomodulatory Properties. Current Pharmaceutical Design 24, 1054–1066. 10.2174/1381612824666180327165012.

13. Nolan, E.M., and Walsh, C.T. (2009). How nature morphs peptide scaffolds into antibiotics. Chembiochem 10, 34–53. 10.1002/cbic.200800438.

14. Singh, N., and Abraham, J. (2014). Ribosomally synthesized peptides from natural sources. J Antibiot 67, 277–289. 10.1038/ja.2013.138.

15. Maasch, J.R.M.A., Torres, M.D.T., Melo, M.C.R., and Fuente-Nunez, C. de la (2022). Molecular de-extinction of ancient antimicrobial peptides enabled by machine learning. bioRxiv 2022.11.15.516443. 10.1101/2022.11.15.516443.

16. García-Bayona, L., and Comstock, L.E. (2018). Bacterial antagonism in host-associated microbial communities. Science 361. 10.1126/science.aat2456.

17. Anderson, M.C., Vonaesch, P., Saffarian, A., Marteyn, B.S., and Sansonetti, P.J. (2017). Shigella sonnei encodes a functional T6SS used for interbacterial competition and niche occupancy. Cell Host Microbe 21. 10.1016/j.chom.2017.05.004.

18. Krismer, B., Weidenmaier, C., Zipperer, A., and Peschel, A. (2017). The commensal lifestyle of Staphylococcus aureus and its interactions with the nasal microbiota. Nat. Rev. Microbiol 15. 10.1038/nrmicro.2017.104.

19. Zhao, W., Caro, F., Robins, W., and Mekalanos, J.J. (2018). Antagonism toward the intestinal microbiota and its effect on Vibrio cholerae virulence. Science 359. 10.1126/science.aap8775.

20. Quereda, J.J. (2017). Listeriolysin S is a streptolysin s-like virulence factor that targets exclusively prokaryotic cells in vivo. mBio 8. 10.1128/mBio.00259-17.

21. Quereda, J.J. (2016). Bacteriocin from epidemic Listeria strains alters the host intestinal microbiota to favor infection. Proc. Natl Acad. Sci. USA 113. 10.1073/pnas.1523899113.

22. Gomes, B., Augusto, M.T., Felício, M.R., Hollmann, A., Franco, O.L., Gonçalves, S., and Santos, N.C. (2018). Designing improved active peptides for therapeutic approaches against infectious diseases. Biotechnol Adv 36, 415–429. 10.1016/j.biotechadv.2018.01.004.

23. Lesiuk, M., Paduszyńska, M., and Greber, K.E. (2022). Synthetic Antimicrobial Immunomodulatory Peptides: Ongoing Studies and Clinical Trials. Antibiotics (Basel) 11, 1062. 10.3390/antibiotics11081062.

24. Mahlapuu, M., Håkansson, J., Ringstad, L., and Björn, C. (2016). Antimicrobial Peptides: An Emerging Category of Therapeutic Agents. Frontiers in Cellular and Infection Microbiology 6.

25. Baquero, F., Lanza, V.F., Baquero, M.R., Campo, R., and Bravo-Vázquez, D.A. (2019). Microcins in Enterobacteriaceae: peptide antimicrobials in the eco-active intestinal chemosphere. Front. Microbiol. 10. 10.3389/fmicb.2019.02261.

26. Kim, S.G. (2019). Microbiota-derived lantibiotic restores resistance against vancomycin-resistant Enterococcus. Nature 572. 10.1038/s41586-019-1501-z.

27. Nakatsuji, T. (2021). Development of a human skin commensal microbe for bacteriotherapy of atopic dermatitis and use in a phase 1 randomized clinical trial. Nat. Med. 27. 10.1038/s41591-021-01256-2.

28. Millette, M., Cornut, G., Dupont, C., Shareck, F., Archambault, D., and Lacroix, M. (2008). Capacity of human nisin-and pediocin-producing lactic Acid bacteria to reduce intestinal colonization by vancomycin-resistant enterococci. Appl Environ Microbiol 74, 1997–2003. 10.1128/AEM.02150-07.

29. O’Reilly, C., O’Connor, P.M., O’Sullivan, Ó., Rea, M.C., Hill, C., and Ross, R.P. (2022). Impact of nisin on Clostridioides difficile and microbiota composition in a faecal fermentation model of the human colon. J Appl Microbiol 132, 1397–1408. 10.1111/jam.15250.

30. Spohn, R., Daruka, L., Lázár, V., Martins, A., Vidovics, F., Grézal, G., Méhi, O., Kintses, B., Számel, M., Jangir, P.K., et al. (2019). Integrated evolutionary analysis reveals antimicrobial peptides with limited resistance. Nat Commun 10, 4538. 10.1038/s41467-019-12364-6.

31. Cesaro, A., Torres, M.D.T., Gaglione, R., Dell’Olmo, E., Di Girolamo, R., Bosso, A., Pizzo, E., Haagsman, H.P., Veldhuizen, E.J.A., de la Fuente-Nunez, C., et al. (2022). Synthetic Antibiotic Derived from Sequences Encrypted in a Protein from Human Plasma. ACS Nano 16, 1880– 1895. 10.1021/acsnano.1c04496.

32. Hyatt, D., Chen, G.-L., LoCascio, P.F., Land, M.L., Larimer, F.W., and Hauser, L.J. (2010). Prodigal: prokaryotic gene recognition and translation initiation site identification. BMC Bioinformatics 11, 119. 10.1186/1471-2105-11-119.

33. Ahrens, C.H., Wade, J.T., Champion, M.M., and Langer, J.D. (2022). A Practical Guide to Small Protein Discovery and Characterization Using Mass Spectrometry. J Bacteriol 204, e0035321. 10.1128/JB.00353-21.

34. Storz, G., Wolf, Y.I., and Ramamurthi, K.S. (2014). Small Proteins Can No Longer Be Ignored. Annu. Rev. Biochem. 83, 753–777. 10.1146/annurev-biochem-070611-102400.

35. Su, M., Ling, Y., Yu, J., Wu, J., and Xiao, J. (2013). Small proteins: untapped area of potential biological importance. Front Genet 4. 10.3389/fgene.2013.00286.

36. Sberro, H., Fremin, B.J., Zlitni, S., Edfors, F., Greenfield, N., Snyder, M.P., Pavlopoulos, G.A., Kyrpides, N.C., and Bhatt, A.S. (2019). Large-Scale Analyses of Human Microbiomes Reveal Thousands of Small, Novel Genes. Cell 178, 1245–1259.e14. 10.1016/j.cell.2019.07.016.

37. Donia, M.S. (2014). A systematic analysis of biosynthetic gene clusters in the human microbiome reveals a common family of antibiotics. Cell 158. 10.1016/j.cell.2014.08.032.

38. Fingerhut, L.C.H.W., Miller, D.J., Strugnell, J.M., Daly, N.L., and Cooke, I.R. (2020). ampir: an R package for fast genome-wide prediction of antimicrobial peptides. Bioinformatics 36, 5262–5263. 10.1093/bioinformatics/btaa653.

39. Sugimoto, Y. (2019). A metagenomic strategy for harnessing the chemical repertoire of the human microbiome. Science 366. 10.1126/science.aax9176.

40. Santos-Júnior, C.D., Pan, S., Zhao, X.-M., and Coelho, L.P. (2020). Macrel: antimicrobial peptide screening in genomes and metagenomes. PeerJ 8, e10555. 10.7717/peerj.10555.

41. Mende, D.R., Letunic, I., Maistrenko, O.M., Schmidt, T.S.B., Milanese, A., Paoli, L., Hernández-Plaza, A., Orakov, A.N., Forslund, S.K., Sunagawa, S., et al. (2020). proGenomes2: an improved database for accurate and consistent habitat, taxonomic and functional annotations of prokaryotic genomes. Nucleic Acids Research 48, D621–D625. 10.1093/nar/gkz1002.

42. Navidinia, M. (2016). The clinical importance of emerging ESKAPE pathogens in nosocomial infections. Archives of Advances in Biosciences 7, 43–57. 10.22037/jps.v7i3.12584.

43. Mulani, M.S., Kamble, E.E., Kumkar, S.N., Tawre, M.S., and Pardesi, K.R. (2019). Emerging Strategies to Combat ESKAPE Pathogens in the Era of Antimicrobial Resistance: A Review. Front Microbiol 10, 539. 10.3389/fmicb.2019.00539.

44. Shi, G., Kang, X., Dong, F., Liu, Y., Zhu, N., Hu, Y., Xu, H., Lao, X., and Zheng, H. (2021). DRAMP 3.0: an enhanced comprehensive data repository of antimicrobial peptides. Nucleic Acids Research. 10.1093/nar/gkab651.

45. Hao, Y., Zhang, L., Niu, Y., Cai, T., Luo, J., He, S., Zhang, B., Zhang, D., Qin, Y., Yang, F., et al. (2018). SmProt: a database of small proteins encoded by annotated coding and non-coding RNA loci. Briefings in Bioinformatics 19, 636–643. 10.1093/bib/bbx005.

46. Coelho, L.P., Alves, R., del Río, Á.R., Myers, P.N., Cantalapiedra, C.P., Giner-Lamia, J., Schmidt, T.S., Mende, D.R., Orakov, A., Letunic, I., et al. (2022). Towards the biogeography of prokaryotic genes. Nature 601, 252–256. 10.1038/s41586-021-04233-4.

47. Murphy, L.R., Wallqvist, A., and Levy, R.M. (2000). Simplified amino acid alphabets for protein fold recognition and implications for folding. Protein Engineering, Design and Selection 13, 149–152. 10.1093/protein/13.3.149.

48. Huerta-Cepas, J., Szklarczyk, D., Heller, D., Hernández-Plaza, A., Forslund, S.K., Cook, H., Mende, D.R., Letunic, I., Rattei, T., Jensen, L.J., et al. (2019). eggNOG 5.0: a hierarchical, functionally and phylogenetically annotated orthology resource based on 5090 organisms and 2502 viruses. Nucleic Acids Research 47, D309–D314. 10.1093/nar/gky1085.

49. Río, Á.R. del Giner-Lamia, J., Cantalapiedra, C.P., Botas, J., Deng, Z., Hernández-Plaza, A., Paoli, L., Schmidt, T.S.B., Sunagawa, S., Bork, P., et al. (2022). Functional and evolutionary significance of unknown genes from uncultivated taxa. bioRxiv 2022.01.26.477801. 10.1101/2022.01.26.477801.

50. Hurtado-Rios, J.J., Carrasco-Navarro, U., Almanza-Pérez, J.C., and Ponce-Alquicira, E. (2022). Ribosomes: The New Role of Ribosomal Proteins as Natural Antimicrobials. Int J Mol Sci 23, 9123. 10.3390/ijms23169123.

51. Blaustein, R.A., McFarland, A.G., Ben Maamar, S., Lopez, A., Castro-Wallace, S., and Hartmann, E.M. (2019). Pangenomic Approach To Understanding Microbial Adaptations within a Model Built Environment, the International Space Station, Relative to Human Hosts and Soil. mSystems 4, e00281–18. 10.1128/mSystems.00281-18.

52. Collins, F.W.J., Mesa-Pereira, B., O’Connor, P.M., Rea, M.C., Hill, C., and Ross, R.P. (2018). Reincarnation of Bacteriocins From the Lactobacillus Pangenomic Graveyard. Front Microbiol 9, 1298. 10.3389/fmicb.2018.01298.

53. Simmons, W.L., Daubenspeck, J.M., Osborne, J.D., Balish, M.F., Waites, K.B., and Dybvig, K. (2013). Type 1 and type 2 strains of Mycoplasma pneumoniae form different biofilms. Microbiology (Reading) 159, 737–747. 10.1099/mic.0.064782-0.

54. Diaz, M.H., Desai, H.P., Morrison, S.S., Benitez, A.J., Wolff, B.J., Caravas, J., Read, T.D., Dean, D., and Winchell, J.M. (2017). Comprehensive bioinformatics analysis of Mycoplasma pneumoniae genomes to investigate underlying population structure and type-specific determinants. Plos One 12, e0174701. 10.1371/journal.pone.0174701.

55. Parks, D.H., Rinke, C., Chuvochina, M., Chaumeil, P.-A., Woodcroft, B.J., Evans, P.N., Hugenholtz, P., and Tyson, G.W. (2017). Recovery of nearly 8,000 metagenome-assembled genomes substantially expands the tree of life. Nature Microbiology 2, 1533–1542. 10.1038/s41564-017-0012-7.

56. Parks, D.H., Chuvochina, M., Chaumeil, P.-A., Rinke, C., Mussig, A.J., and Hugenholtz, P. (2020). A complete domain-to-species taxonomy for Bacteria and Archaea. Nature Biotechnology, 1–8. 10.1038/s41587-020-0501-8.

57. Valles-Colomer, M., Blanco-Míguez, A., Manghi, P., Asnicar, F., Dubois, L., Golzato, D., Armanini, F., Cumbo, F., Huang, K.D., Manara, S., et al. (2023). The person-to-person transmission landscape of the gut and oral microbiomes. Nature 614, 125–135. 10.1038/s41586-022-05620-1.

58. Pirtskhalava, M., Amstrong, A.A., Grigolava, M., Chubinidze, M., Alimbarashvili, E., Vishnepolsky, B., Gabrielian, A., Rosenthal, A., Hurt, D.E., and Tartakovsky, M. (2021). DBAASP v3: database of antimicrobial/cytotoxic activity and structure of peptides as a resource for development of new therapeutics. Nucleic Acids Research 49, D288–D297. 10.1093/nar/gkaa991.

59. Wang, G., Li, X., and Wang, Z. (2016). APD3: the antimicrobial peptide database as a tool for research and education. Nucleic Acids Res 44, D1087–1093. 10.1093/nar/gkv1278.

60. Torres, M.D.T., Sothiselvam, S., Lu, T.K., and de la Fuente-Nunez, C. (2019). Peptide Design Principles for Antimicrobial Applications. J Mol Biol 431, 3547–3567. 10.1016/j.jmb.2018.12.015.

61. Lifson, S., and Sander, C. (1979). Antiparallel and parallel β-strands differ in amino acid residue preferences. Nature 282, 109–111. 10.1038/282109a0.

62. Cullen, T.W., Schofield, W.B., Barry, N.A., Putnam, E.E., Rundell, E.A., Trent, M.S., Degnan, P.H., Booth, C.J., Yu, H., and Goodman, A.L. (2015). Antimicrobial peptide resistance mediates resilience of prominent gut commensals during inflammation. Science 347, 170–175. 10.1126/science.1260580.

63. Torres, M.D.T., Pedron, C.N., Araújo, I., Silva Jr., P.I., Silva, F.D., and Oliveira, V.X. (2017). Decoralin Analogs with Increased Resistance to Degradation and Lower Hemolytic Activity. ChemistrySelect 2, 18–23. 10.1002/slct.201601590.

64. Torres, M.D.T., Pedron, C.N., Higashikuni, Y., Kramer, R.M., Cardoso, M.H., Oshiro, K.G.N., Franco, O.L., Silva Junior, P.I., Silva, F.D., Oliveira Junior, V.X., et al. (2018). Structure-function-guided exploration of the antimicrobial peptide polybia-CP identifies activity determinants and generates synthetic therapeutic candidates. Commun Biol 1, 1–16. 10.1038/s42003-018-0224-2.

65. Silva, O.N., Torres, M.D.T., Cao, J., Alves, E.S.F., Rodrigues, L.V., Resende, J.M., Lião, L.M., Porto, W.F., Fensterseifer, I.C.M., Lu, T.K., et al. (2020). Repurposing a peptide toxin from wasp venom into antiinfectives with dual antimicrobial and immunomodulatory properties. Proc Natl Acad Sci U S A 117, 26936–26945. 10.1073/pnas.2012379117.

66. Morris, F.C., Dexter, C., Kostoulias, X., Uddin, M.I., and Peleg, A.Y. (2019). The Mechanisms of Disease Caused by Acinetobacter baumannii. Frontiers in Microbiology 10.

67. Petruschke, H., Schori, C., Canzler, S., Riesbeck, S., Poehlein, A., Daniel, R., Frei, D., Segessemann, T., Zimmerman, J., Marinos, G., et al. (2021). Discovery of novel community-relevant small proteins in a simplified human intestinal microbiome. Microbiome 9, 55. 10.1186/s40168-020-00981-z.

68. Galzitskaya, O.V. (2021). Exploring Amyloidogenicity of Peptides From Ribosomal S1 Protein to Develop Novel AMPs. Front Mol Biosci 8, 705069. 10.3389/fmolb.2021.705069.

69. Lazzaro, B.P., Zasloff, M., and Rolff, J. (2020). Antimicrobial peptides: Application informed by evolution. Science 368, eaau5480. 10.1126/science.aau5480.

70. Gebhard, S. (2012). ABC transporters of antimicrobial peptides in Firmicutes bacteria - phylogeny, function and regulation. Mol Microbiol 86, 1295–1317. 10.1111/mmi.12078.

71. Claesen, J., Spagnolo, J.B., Ramos, S.F., Kurita, K.L., Byrd, A.L., Aksenov, A.A., Melnik, A.V., Wong, W.R., Wang, S., Hernandez, R.D., et al. (2020). A Cutibacterium acnes antibiotic modulates human skin microbiota composition in hair follicles. Sci Transl Med 12, eaay5445. 10.1126/scitranslmed.aay5445.

72. Coelho, L.P., Alves, R., Monteiro, P., Huerta-Cepas, J., Freitas, A.T., and Bork, P. (2019). NG-meta-profiler: fast processing of metagenomes using NGLess, a domain-specific language. Microbiome 7, 84. 10.1186/s40168-019-0684-8.

73. Coelho, L.P. (2017). Jug: Software for Parallel Reproducible Computation in Python. Journal of Open Research Software 5, 30. 10.5334/jors.161.

74. Fu, L., Niu, B., Zhu, Z., Wu, S., and Li, W. (2012). CD-HIT: accelerated for clustering the next-generation sequencing data. Bioinformatics 28, 3150–3152. 10.1093/bioinformatics/bts565.

75. Steinegger, M., and Söding, J. (2017). MMseqs2 enables sensitive protein sequence searching for the analysis of massive data sets. Nature Biotechnology 35, 1026–1028. 10.1038/nbt.3988.

76. Van Rossum, G. (2020). Python Release Python 3.8.2. Python.org. https://www.python.org/downloads/release/python-382/.

77. Hunter, J.D. (2007). Matplotlib: A 2D Graphics Environment. Computing in Science Engineering 9, 90–95. 10.1109/MCSE.2007.55.

78. Harris, C.R., Millman, K.J., van der Walt, S.J., Gommers, R., Virtanen, P., Cournapeau, D., Wieser, E., Taylor, J., Berg, S., Smith, N.J., et al. (2020). Array programming with NumPy. Nature 585, 357–362. 10.1038/s41586-020-2649-2.

79. McKinney, W. (2010). Data Structures for Statistical Computing in Python. Proceedings of the 9th Python in Science Conference, 56–61. 10.25080/Majora-92bf1922-00a.

80. Virtanen, P., Gommers, R., Oliphant, T.E., Haberland, M., Reddy, T., Cournapeau, D., Burovski, E., Peterson, P., Weckesser, W., Bright, J., et al. (2020). SciPy 1.0: fundamental algorithms for scientific computing in Python. Nat Methods 17, 261–272. 10.1038/s41592-019-0686-2.

81. Pedregosa, F., Varoquaux, G., Gramfort, A., Michel, V., Thirion, B., Grisel, O., Blondel, M., Prettenhofer, P., Weiss, R., Dubourg, V., et al. (2011). Scikit-learn: Machine Learning in Python. Machine Learning in Python 12, 2825–2830.

82. The scikit-bio development team (2020). scikit-bio: A Bioinformatics Library for Data Scientists, Students, and Developers. http://scikit-bio.org/

83. Cock, P.J.A., Antao, T., Chang, J.T., Chapman, B.A., Cox, C.J., Dalke, A., Friedberg, I., Hamelryck, T., Kauff, F., Wilczynski, B., et al. (2009). Biopython: freely available Python tools for computational molecular biology and bioinformatics. Bioinformatics 25, 1422–1423. 10.1093/bioinformatics/btp163.

84. Cantalapiedra, C.P., Hernández-Plaza, A., Letunic, I., Bork, P., and Huerta-Cepas, J. (2021). eggNOG-mapper v2: Functional Annotation, Orthology Assignments, and Domain Prediction at the Metagenomic Scale. Molecular Biology and Evolution 38, 5825–5829. 10.1101/2021.06.03.446934.

85. Eddy, S.R. (2011). Accelerated Profile HMM Searches. PLoS Computational Biology 7, e1002195. 10.1371/journal.pcbi.1002195.

86. Müller, A.T., Gabernet, G., Hiss, J.A., and Schneider, G. (2017). modlAMP: Python for antimicrobial peptides. Bioinformatics 33, 2753–2755. 10.1093/bioinformatics/btx285.

87. Price, M.N., Dehal, P.S., and Arkin, A.P. (2010). FastTree 2 – Approximately Maximum-Likelihood Trees for Large Alignments. PLOS ONE 5, e9490. 10.1371/journal.pone.0009490.

88. Jain, C., Rodriguez-R, L.M., Phillippy, A.M., Konstantinidis, K.T., and Aluru, S. (2017). High-throughput ANI Analysis of 90K Prokaryotic Genomes Reveals Clear Species Boundaries. bioRxiv, 225342. 10.1101/225342.

89. Li, D., Luo, R., Liu, C.M., Leung, C.M., Ting, H.F., Sadakane, K., Yamashita, H., and Lam, T.W. (2016). MEGAHIT v1.0: A fast and scalable metagenome assembler driven by advanced methodologies and community practices. 102, 3–11. 10.1016/j.ymeth.2016.02.020.

90. The UniProt Consortium (2021). UniProt: the universal protein knowledgebase in 2021. Nucleic Acids Research 49, D480–D489. 10.1093/nar/gkaa1100.

91. Aguilera-Mendoza, L., Marrero-Ponce, Y., Beltran, J.A., Tellez Ibarra, R., Guillen-Ramirez, H.A., and Brizuela, C.A. (2019). Graph-based data integration from bioactive peptide databases of pharmaceutical interest: toward an organized collection enabling visual network analysis. Bioinformatics 35, 4739–4747. 10.1093/bioinformatics/btz260.

92. Mistry, J., Chuguransky, S., Williams, L., Qureshi, M., Salazar, G.A., Sonnhammer, E.L.L., Tosatto, S.C.E., Paladin, L., Raj, S., Richardson, L.J., et al. (2021). Pfam: The protein families database in 2021. Nucleic Acids Research 49, D412–D419. 10.1093/nar/gkaa913.

93. Eberhardt, R.Y., Haft, D.H., Punta, M., Martin, M., O’Donovan, C., and Bateman, A. (2012). AntiFam: a tool to help identify spurious ORFs in protein annotation. Database (Oxford) 2012, bas003. 10.1093/database/bas003.

94. NCBI Resource Coordinators (2015). Database resources of the National Center for Biotechnology Information. Nucleic Acids Research 43, D6–D17. 10.1093/nar/gku1130.

95. Courtot, M., Cherubin, L., Faulconbridge, A., Vaughan, D., Green, M., Richardson, D., Harrison, P., Whetzel, P.L., Parkinson, H., and Burdett, T. (2019). BioSamples database: an updated sample metadata hub. Nucleic Acids Research 47, D1172–D1178. 10.1093/nar/gky1061.

96. Mirdita, M., Steinegger, M., Breitwieser, F., Söding, J., and Levy Karin, E. (2021). Fast and sensitive taxonomic assignment to metagenomic contigs. Bioinformatics. 10.1093/bioinformatics/btab184.

97. Finn, R.D., Coggill, P., Eberhardt, R.Y., Eddy, S.R., Mistry, J., Mitchell, A.L., Potter, S.C., Punta, M., Qureshi, M., Sangrador-Vegas, A., et al. (2016). The Pfam protein families database: towards a more sustainable future. Nucleic acids research 44, D279–85. 10.1093/nar/gkv1344.

98. Solis, A.D. (2015). Amino acid alphabet reduction preserves fold information contained in contact interactions in proteins. Proteins: Structure, Function, and Bioinformatics 83, 2198–2216. 10.1002/prot.24936.

99. Peterson, E.L., Kondev, J., Theriot, J.A., and Phillips, R. (2009). Reduced amino acid alphabets exhibit an improved sensitivity and selectivity in fold assignment. Bioinformatics 25, 1356– 1362. 10.1093/bioinformatics/btp164.

100. Smith, T.F., and Waterman, M.S. (1981). Identification of Common Molecular Subsequences. J. Mol. Biol. 147, 195–197. 10.1016/0022-2836(81)90087-5.

101. Karlin, S., and Altschul, S.F. (1990). Methods for assessing the statistical significance of molecular sequence features by using general scoring schemes. Proc Natl Acad Sci U S A 87, 2264–2268. 10.1073/pnas.87.6.2264.

102. Altschul, S., Madden, T., Schaffer, A., Zhang, J., Zhang, Z., Miller, W., and Lipman, D. (1997). Gapped BLAST and PSI-BLAST: a new generation of protein database search programs. Nucleic Acids Res. 25, 3389–3402.

103. Camacho, C., Coulouris, G., Avagyan, V., Ma, N., Papadopoulos, J., Bealer, K., and Madden, T.L. (2009). BLAST+: architecture and applications. BMC Bioinformatics 10, 421. 10.1186/1471-2105-10-421.

104. Washietl, S., Findeiß, S., Müller, S.A., Kalkhof, S., Bergen, M. von Hofacker, I.L., Stadler, P.F., and Goldman, N. (2011). RNAcode: Robust discrimination of coding and noncoding regions in comparative sequence data. RNA 17, 578–594. 10.1261/rna.2536111.

105. Li, H., and Durbin, R. (2009). Fast and accurate short read alignment with Burrows-Wheeler transform. Bioinformatics 25, 1754–1760. 10.1093/bioinformatics/btp324.

106. Seabold, S., and Perktold, J. (2010). Statsmodels: Econometric and Statistical Modeling with Python. Proceedings of the 9th Python in Science Conference, 92–96. 10.25080/Majora-92bf1922-011.

107. Milanese, A., Mende, D.R., Paoli, L., Salazar, G., Ruscheweyh, H.-J., Cuenca, M., Hingamp, P., Alves, R., Costea, P.I., Coelho, L.P., et al. (2019). Microbial abundance, activity and population genomic profiling with mOTUs2. Nat Commun 10, 1014. 10.1038/s41467-019-08844-4.

108. Sélem-Mojica, N., Aguilar, C., Gutiérrez-García, K., Martínez-Guerrero, C.E., and Barona-Gómez, F. (2019). EvoMining reveals the origin and fate of natural product biosynthetic enzymes. Microb Genom 5, e000260. 10.1099/mgen.0.000260.

109. Rodriguez-R, L.M., Conrad, R.E., Feistel, D.J., Viver, T., Rosselló-Móra, R., and Konstantinidis, K.T. (2022). A natural definition for a bacterial strain and clonal complex. 2022.06.27.497766. 10.1101/2022.06.27.497766.

110. Li, H., Handsaker, B., Wysoker, A., Fennell, T., Ruan, J., Homer, N., Marth, G., Abecasis, G., Durbin, R., and 1000 Genome Project Data Processing Subgroup (2009). The Sequence Alignment/Map format and SAMtools. Bioinformatics 25, 2078–2079. 10.1093/bioinformatics/btp352.

111. Quinlan, A.R., and Hall, I.M. (2010). BEDTools: A flexible suite of utilities for comparing genomic features. Bioinformatics 26, 841–842. 10.1093/bioinformatics/btq033.

112. Sievers, F., Wilm, A., Dineen, D., Gibson, T.J., Karplus, K., Li, W., Lopez, R., McWilliam, H., Remmert, M., Söding, J., et al. (2011). Fast, scalable generation of high-quality protein multiple sequence alignments using Clustal Omega. Mol Syst Biol 7, 539. 10.1038/msb.2011.75.

113. Venturini, E., Svensson, S.L., Maaß, S., Gelhausen, R., Eggenhofer, F., Li, L., Cain, A.K., Parkhill, J., Becher, D., Backofen, R., et al. (2020). A global data-driven census of Salmonella small proteins and their potential functions in bacterial virulence. microLife 1, uqaa002. 10.1093/femsml/uqaa002.

114. Kosakovsky Pond, S.L., Poon, A.F.Y., Velazquez, R., Weaver, S., Hepler, N.L., Murrell, B., Shank, S.D., Magalis, B.R., Bouvier, D., Nekrutenko, A., et al. (2019). HyPhy 2.5—A Customizable Platform for Evolutionary Hypothesis Testing Using Phylogenies. Mol Biol Evol 37, 295–299. 10.1093/molbev/msz197.

115. Murrell, B., Weaver, S., Smith, M.D., Wertheim, J.O., Murrell, S., Aylward, A., Eren, K., Pollner, T., Martin, D.P., Smith, D.M., et al. (2015). Gene-Wide Identification of Episodic Selection. Molecular Biology and Evolution 32, 1365–1371. 10.1093/molbev/msv035.

116. Buchfink, B., Xie, C., and Huson, D.H. (2014). Fast and sensitive protein alignment using DIAMOND. Nature Methods 12, 59–60. 10.1038/nmeth.3176.

117. Kanehisa, M., and Sato, Y. (2020). KEGG Mapper for inferring cellular functions from protein sequences. Protein Sci. 29, 28–35. 10.1002/pro.3711.

118. Alcock, B.P., Raphenya, A.R., Lau, T.T.Y., Tsang, K.K., Bouchard, M., Edalatmand, A., Huynh, W., Nguyen, A.-L.V., Cheng, A.A., Liu, S., et al. (2020). CARD 2020: antibiotic resistome surveillance with the comprehensive antibiotic resistance database. Nucleic Acids Res 48, D517– D525. 10.1093/nar/gkz935.

119. Ochoa, R., and Cossio, P. (2021). PepFun: Open Source Protocols for Peptide-Related Computational Analysis. Molecules 26, 1664. 10.3390/molecules26061664.

120. Veltri, D., Kamath, U., and Shehu, A. (2018). Deep learning improves antimicrobial peptide recognition. Bioinformatics 34, 2740–2747. 10.1093/bioinformatics/bty179.

121. Lawrence, T.J., Carper, D.L., Spangler, M.K., Carrell, A.A., Rush, T.A., Minter, S.J., Weston, D.J., and Labbé, J.L. (2021). amPEPpy 1.0: a portable and accurate antimicrobial peptide prediction tool. Bioinformatics 37, 2058–2060. 10.1093/bioinformatics/btaa917.

122. Su, X., Xu, J., Yin, Y., Quan, X., and Zhang, H. (2019). Antimicrobial peptide identification using multi-scale convolutional network. BMC Bioinformatics 20, 730. 10.1186/s12859-019-3327-y.

123. Li, C., Sutherland, D., Hammond, S.A., Yang, C., Taho, F., Bergman, L., Houston, S., Warren, R.L., Wong, T., Hoang, L.M.N., et al. (2022). AMPlify: attentive deep learning model for discovery of novel antimicrobial peptides effective against WHO priority pathogens. BMC Genomics 23, 77. 10.1186/s12864-022-08310-4.

124. Kochendoerfer, G.G., and Kent, S.B. (1999). Chemical protein synthesis. Curr Opin Chem Biol 3, 665–671. 10.1016/s1367-5931(99)00024-1.

125. Sheppard, R. (2003). The fluorenylmethoxycarbonyl group in solid phase synthesis. J Pept Sci 9, 545–552. 10.1002/psc.479.

126. Palomo, J.M. (2014). Solid-phase peptide synthesis: an overview focused on the preparation of biologically relevant peptides. RSC Adv. 4, 32658–32672. 10.1039/C4RA02458C.

127. Behrendt, R., White, P., and Offer, J. (2016). Advances in Fmoc solid-phase peptide synthesis. Journal of Peptide Science 22, 4–27. 10.1002/psc.2836.

128. Schmidt, T.S.B., Li, S.S., Maistrenko, O.M., Akanni, W., Coelho, L.P., Dolai, S., Fullam, A., Glazek, A.M., Hercog, R., Herrema, H., et al. (2022). Drivers and determinants of strain dynamics following fecal microbiota transplantation. Nat Med 28, 1902–1912. 10.1038/s41591-022-01913-0.

129. Wiegand, I., Hilpert, K., and Hancock, R.E.W. (2008). Agar and broth dilution methods to determine the minimal inhibitory concentration (MIC) of antimicrobial substances. Nat Protoc 3, 163–175. 10.1038/nprot.2007.521.

130. Micsonai, A., Moussong, É., Wien, F., Boros, E., Vadászi, H., Murvai, N., Lee, Y.-H., Molnár, T., Réfrégiers, M., Goto, Y., et al. (2022). BeStSel: webserver for secondary structure and fold prediction for protein CD spectroscopy. Nucleic Acids Research 50, W90–W98. 10.1093/nar/gkac345.

131. Santos-Júnior, C.D., Schmidt, T.S.B., Fullam, A., Duan, Y., Bork, P., Zhao, X.-M., and Coelho, L.P. (2021). AMPSphere : the worldwide survey of prokaryotic antimicrobial peptides. Zenodo (Database). 10.5281/zenodo.4606582.

132. Heintz-Buschart, A., May, P., Laczny, C.C., Lebrun, L.A., Bellora, C., Krishna, A., Wampach, L., Schneider, J.G., Hogan, A., de Beaufort, C., et al. (2016). Integrated multi-omics of the human gut microbiome in a case study of familial type 1 diabetes. Nat Microbiol 2, 16180. 10.1038/nmicrobiol.2016.180.

## Supplemental references

S1 Santos-Júnior CD, Pan S, Zhao X-M, Coelho LP. Macrel: antimicrobial peptide screening in genomes and metagenomes. PeerJ 2020; 8:e10555.

S2. Shi G, Kang X, Dong F, et al. DRAMP 3.0: an enhanced comprehensive data repository of antimicrobial peptides. Nucleic Acids Research 2021.

S3. Coelho LP, Alves R, del Río ÁR, et al. Towards the biogeography of prokaryotic genes. Nature 2022; 601:252–256.

S4. Huerta-Cepas J, Szklarczyk D, Heller D, et al. eggNOG 5.0: a hierarchical, functionally and phylogenetically annotated orthology resource based on 5090 organisms and 2502 viruses. Nucleic Acids Research 2019; 47:D309–D314.

S5. Kanehisa M, Sato Y. KEGG Mapper for inferring cellular functions from protein sequences. Protein Sci 2020; 29:28–35.

S6. Alcock BP, Raphenya AR, Lau TTY, et al. CARD 2020: antibiotic resistome surveillance with the comprehensive antibiotic resistance database. Nucleic Acids Res 2020; 48:D517–D525.

S7. Cantalapiedra CP, Hernández-Plaza A, Letunic I, Bork P, Huerta-Cepas J. eggNOG-mapper v2: Functional Annotation, Orthology Assignments, and Domain Prediction at the Metagenomic Scale.; 2021:2021.06.03.446934.

